# A transmission chain linking *Mycobacterium ulcerans* with *Aedes notoscriptus* mosquitoes, possums and human Buruli ulcer cases in southeastern Australia

**DOI:** 10.1101/2023.05.07.539718

**Authors:** Peter T. Mee, Andrew H. Buultjens, Jane Oliver, Karen Brown, Jodie C. Crowder, Jessica L. Porter, Emma C. Hobbs, Louise M. Judd, George Taiaroa, Natsuda Puttharak, Deborah A. Williamson, Kim R. Blasdell, Ee Laine Tay, Rebecca Feldman, Mutizwa Odwell Muzari, Chris Sanders, Stuart Larsen, Simon R. Crouch, Paul D. R. Johnson, John R. Wallace, David J. Price, Ary A. Hoffmann, Katherine B. Gibney, Timothy P. Stinear, Stacey E. Lynch

## Abstract

In temperate southeastern Australia over the past two decades there has been a marked progressive increase in human cases of Buruli ulcer, an infection of subcutaneous tissue caused by *Mycobacterium ulcerans*. Native possums are the major local environmental reservoir of *M. ulcerans* as they not only develop Buruli lesions but they also shed *M. ulcerans* in their excreta. However the way humans acquire *M. ulcerans* from possums has not been determined. Previous case-control studies, insect field surveys and vector competence studies have suggested a role for mosquitoes in *M. ulcerans* transmission between possums and humans. To explore these links we conducted an extensive, 4-month structured mosquito field survey and four *ad hoc* field surveys across an area of 350km^2^ on the Mornington Peninsula, an area endemic for Buruli ulcer to the south of the major metropolitan city of Melbourne. We then compared spatial and temporal patterns of *M. ulcerans*-positive mosquito occurrence with *M. ulcerans*-positive possums (established by previous possum excreta surveys) and human Buruli ulcer cases across the region. We used metabarcoding to assess mosquito blood-feeding host preference and to reconstruct *M. ulcerans* genomes from positive mosquitoes to test epidemiological inferences. We collected 66,325 mosquitoes spanning 26 different species from 180 repeatedly sampled traps over a 4-month period. *Culex molestus* and *Aedes notoscriptus* were the dominant species (42% and 35% of trapped mosquitoes, respectively). PCR screening 25% of trapped mosquitoes revealed a significant association between *M. ulcerans* and *Ae. notoscriptus* (p<0.0001) with a maximum likelihood estimate (MLE) of 5.88 *M. ulcerans* positive mosquitoes per 1,000 tested. Using spatial scanning statistics, we also observed significant overlap between clusters of *M. ulcerans*-positive *Ae. notoscriptus*, *M. ulcerans*-positive possum excreta and human Buruli ulcer cases. Metabarcoding analyses of blood-fed *Ae. notoscriptus* showed individual mosquitoes had fed both on humans and native possums. Enrichment genome sequencing from PCR-positive mosquitoes confirmed shared *M. ulcerans* genome single-nucleotide polymorphism (SNP) profiles between mosquitoes, possum excreta and clinical human isolates within the same regions. These findings indicate that *Ae. notoscriptus* likely transmit *M. ulcerans* in southeastern Australia and highlight mosquito control as a plausible means to control the Buruli ulcer epidemic in our region.

## Introduction

*Mycobacterium ulcerans* is the causative agent of a neglected tropical skin disease called Buruli ulcer, a necrotising infection of skin and subcutaneous tissue ^1^. Buruli ulcer is rarely a fatal condition but can cause severe tissue destruction if not diagnosed and managed effectively ^2^. Buruli ulcer has been described in more than 32 countries worldwide ^3^ and is an ongoing public health issue in west and central Africa ^4^. Buruli ulcer has also been unexpectedly surging in temperate southeastern Australia (Figure 1) and encroaching on the major metropolitan centres of Melbourne (population 5.1 million) and Geelong (population 274,000), with >250 cases routinely notified each year since 2017 to the Victorian State Government Department of Health5.

**Figure 1.**
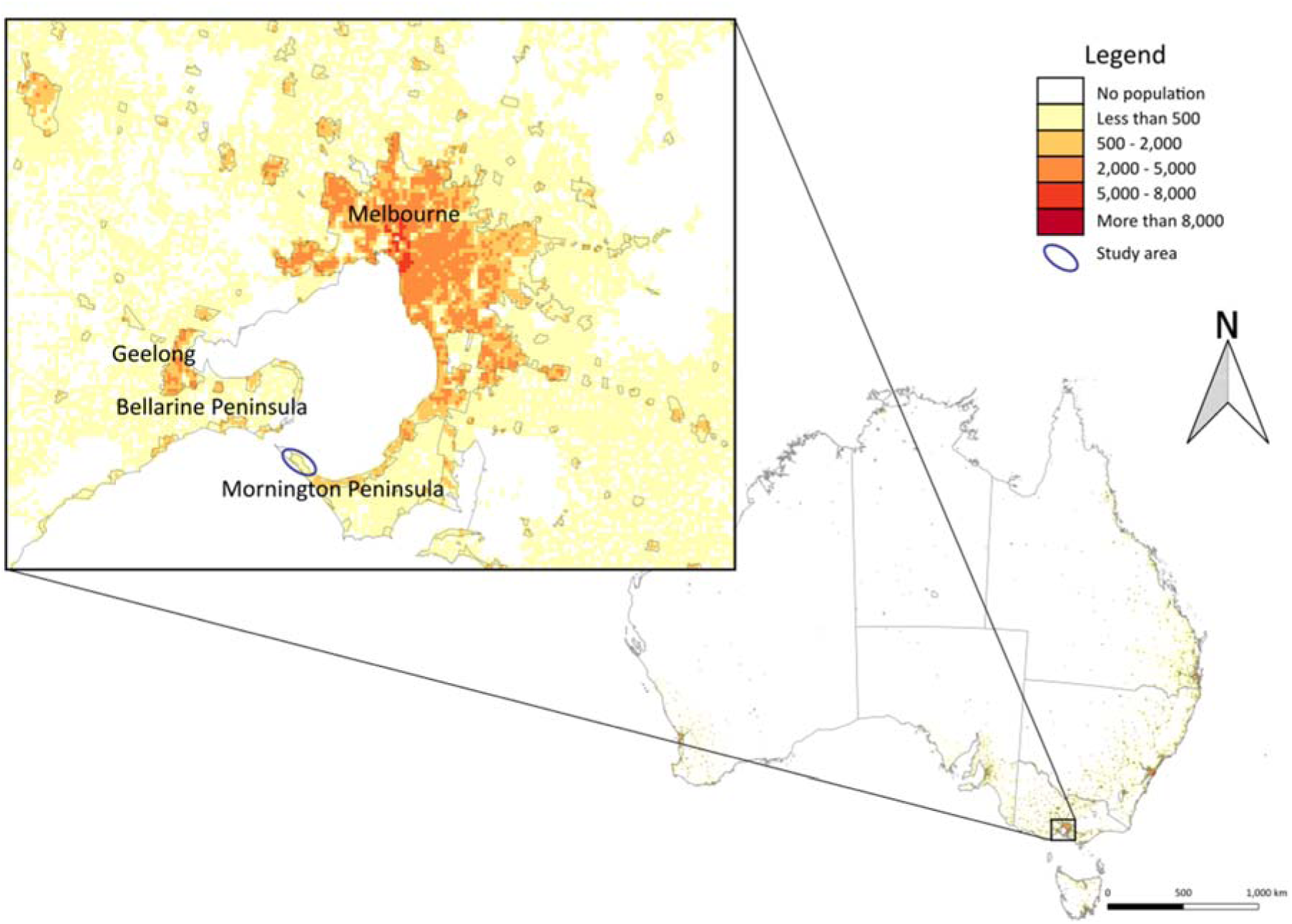
Location of the Mornington Peninsula and the Bellarine Peninsula of Victoria, Australia. Population density is represented on the map from low to high (white to red) based on the human population per square kilometre. The study area is shown within an elipse (inset).

How humans contract Buruli ulcer is a central question that has confounded public health control efforts and intrigued scientists since the discovery of *M. ulcerans* from patients in the Bairnsdale region of Australia in the 1930s and across Africa shortly after ^1,6,7^. Buruli ulcer epidemiology can be unpredictable with a 4-5 month median incubation period and outbreaks emerging in specific geographical areas and then disappearing over a number of years. It is also very challenging to isolate the bacterium in pure culture from the environment, presumably due to its very slow growth, although it can be isolated from human skin lesions. These factors combined have made it incredibly challenging to establish how *M. ulcerans* is spread to humans, despite global efforts to investigate this over more than 80 years ^8^.

The discovery that Buruli ulcer is a zoonosis and that Australian native possums are a major wildlife source of *M. ulcerans* that is intimately linked with disease transmission has addressed one key component of the transmission enigma ^9–14^. The first indications that mosquitoes might be vectors of *M. ulcerans* from possums to humans in Australia came from a series of entomological field surveys in the southeast of the country in response to an increase in Buruli ulcer cases in the seaside township of Point Lonsdale, located on the Bellarine Peninsula ^15^. Among the 12 species identified from a trapping effort that collected 11,500 mosquitoes, five different species were IS2404 PCR positive for *M. ulcerans,* including *Ae. camptorhynchus*, *Ae. notoscriptus*, *Coquillettidia linealis*, *Culex australicus*, and *Anopheles annulipes* (maximum likelihood estimate (MLE) was 4.11/1000 mosquitoes) ^15^. IS2404 is a *M. ulcerans*-specific insertion sequence and molecular target for the gold-standard diagnostic PCR for Buruli ulcer ^16^.

A concurrent case control study performed in the same geographic area identified only two factors associated with the odds of being diagnosed with Buruli ulcer: insect repellent use reduced risk (OR 0.37, 95% CI 0.19–0.69) and being bitten by mosquitoes on the lower legs increased risk (OR 2.60, 95% CI 1.22– 5.53). A variety of outdoor activities were also surveyed but were not independently predictive suggesting that mosquito exposure specifically rather than environmental exposure generally might be the main mode of MU transmission to humans ^17^. In Africa, two case control studies conducted in Cameroon both found use of bed nets as a protective factor against Buruli ulcer; (OR 0.4, 95% CI 0.2-0.9, p=0.04) ^18^ and (OR 0.1, 95% CI 0.03-0.3, p<0.001) ^19^.

However, case control studies and entomological surveys alone are insufficient to indict biological agents as vectors of pathogens. There are formal frameworks used in biomedicine, such as the Barnett criteria ^3^ that build hierarchies of evidence to implicate a candidate disease vector. Here, we build on this aforementioned research to formally address the Barnett criteria, and test the hypothesis that mosquitoes vector *M. ulcerans* to humans ^20–22^. A summary of our findings is shown in Table 1, including the new data presented in this study that specifically address criteria 1-3.

**Table 1:** Summary of evidence to indict mosquitoes as vectors of *M. ulcerans*.

| Barnett Criterion | Evidence | Reference |
| --- | --- | --- |
| 1. Acquire from source reservoir | *M. ulcerans* genome sequences from possum excreta matches genomes recovered from mosquitoes in same area | This study |
| 2. Close links with infected hosts | *M. ulcerans* genome sequences from humans matches genome recovered from mosquitoes in same area. Mosquito bloodmeal analysis confirms human-possum link. | This study |
| 3. Repeat detections from candidate vector | *M. ulcerans* detected with mosquitoes in seven separate field surveys | <sup>15,21</sup>, this study |
| 4. Experimental demonstration of transmission | Mosquito-*M. ulcerans* mechanical transmission demonstrated using murine contaminated skin model | <sup>22</sup> |

In the research presented here, based on a very substantial field survey of >65,000 mosquitoes undertaken over four months in 2019 and 2020, and four smaller *ad hoc* surveys conducted between 2016 and 2021 along the Mornington Peninsula. The study area encompasses an area of 350 km^2^ and is located 90 km south of Melbourne, the capital city of Victoria ^10^ (Figure 1). The area was originally covered in low-lying coastal vegetation ^10^, receives an average annual rainfall of 740 mm, and sits at an average elevation of 60 metres above sea-level ^24^. The Mornington Peninsula region continues to maintain among the highest incidences of Buruli ulcer in the world ^25^ with a conservative local incidence estimate of 55 human cases/100,000 population in 2022 ^5^ ^26^. Here, we employed spatial clustering analyses, bacterial enrichment genome sequencing and mosquito blood meal metabarcoding to continue building a hierarchy of evidence that supports mosquitoes as vectors of *M. ulcerans* from local wildlife reservoirs to humans.

## Material and methods

### Study Site

Insects were collected from the Mornington Peninsula suburbs of Rye (population 8,416), Blairgowrie (population 2,313), Tootgarook (population 2,869) and Capel Sound (population 4,930) ^23^ (Figure 1).

### Arthropod collection and identification

Mosquito trapping campaigns used Biogent Sentinal (BGS) traps (Biogent) that were baited with dry ice pellets to provide a source of CO2 over an intended 12-hour period. BGS traps were set out at dusk and collected at dawn in shaded locations on the grassed edges of the roadways in the study area. GPS locations for all traps were recorded with data collection managed using *Atlas of Medical Entomology* (Gaia Resources, V3.4.4). Trapped mosquitoes were knocked down with CO2 by placing the catch-bag in dry ice before being transported back to the laboratory and kept at −20°C until processing. Mosquito species were morphologically identified using a stereo dissecting microscope (Nikon, SMZ800N) and reference to taxonomic keys ^27–29^.

To assess the potential presence of *M. ulcerans* beyond mosquitoes, other arthropods were collected using Yellow Sticky Traps (YST) and Sticky Ovitraps (SO). Two YST and SO were placed in residential properties where householders had previously noted insect activity to the researchers ^9^. The SO traps were placed on the ground and had hay grass infusion (3 Jack Rabbit (clover/lucerne) pellets (Laucke Mills, Barossa Valley, SA, Australia) in 500 mL water) added to them, while the YST (Bugs for Bugs, Toowoomba, QLD, Australia) were placed on the ground with a 14 cm plant tag plastic T-support (Garden City Plastics, Dandenong, VIC, Australia). Within three to four days of being set, residents were asked to pack up the YST and SO by covering the sticky card with a plastic film, and return the sealed traps to the laboratory, where they were stored at −20°C. Non-mosquito arthropods were morphologically identified to family level and, if PCR positive for *M. ulcerans,* were DNA barcoded for species confirmation by targeting the cytochrome c oxidase subunit I (COI) gene ^30^.

### DNA extraction from mosquitoes

Mosquitoes were sorted by species per trap and by sex and then pooled in 2 mL o-ring tubes, with a maximum of 15 individuals in each pool. A subset of *Ae. notoscriptus* mosquitoes were also screened individually. Mosquitoe(s) were homogenised with 10 x 1.0 mm zirconia-silica beads (BioSpec Products), with 597 µL of Buffer RLT and 2.8 µL of carrier RNA. Homogenisation was performed using a TissueLyser II (Qiagen) at 30 oscillations/sec for 100 sec, repeated twice. Tubes were then centrifuged at 16,000 g for 3 mins. A 550 µL volume of supernatant was transferred into a 96 well deep well plate, with extraction performed as is the protocol for the BioSprint 96 One-For-All Vet Kit (Qiagen). Every eleventh of twelve wells in a 96-well plate was a blank DNA extraction control (seven in total) and a synthetic IS2404 positive control was spiked into one of these seven wells to act as a positive extraction control. Extraction was performed on a KingFisher^TM^ Flex Magnetic Particle Processors (Thermo Scientific).

### DNA extraction from arthropods

Arthropods other than mosquitoes collected on YST or SO were removed from the sticky cards and placed in 1.5 mL microtubes (Eppendorf). Insects were separated by family and trap location, and were pooled with a maximum of 10 individuals from each family per 1.5 mL tube. Samples were extracted non-destructively to allow species confirmation if positive detections occurred. DNA was extracted using the ISOLATE II Genomic DNA Kit (Bioline). Briefly, 25 µL of Proteinase K and 180 µL of Lysis Buffer GL was added to each tube with samples incubated overnight at 56°C. Following incubation, the insects were removed and stored to allow for further morphological identification if required, with the DNA extraction completed on the incubation solution as per manufacturer instructions.

### Synthetic PCR positive control

A synthetic PCR positive control DNA molecule was designed to discriminate false positives due to contamination with positive control DNA versus the authentic IS2404 amplicon. The synthetic positive control was designed to have an amplicon size of 120 bp to easily differentiate from a true IS2404 PCR positive (59 bp) ^16^. The synthetic positive was added at the DNA extraction stage on all 96-well plates, as a positive control for this step and for the subsequent qPCR. The additional DNA sequence used to construct the synthetic positive control was randomly selected from a DNA sequence unlikely to be in the laboratory, in this case, Irrawaddy Dolphin (MK032252).

The synthetic positive control had the sequence: 5’ – TCCT<u>AAAGCACCACGCAGCATCT</u>ATCGCGAGCTTAATCACCATGCCG<u>CGTCCAACGCGATC</u>CCCGCTCGGCAGGGATC CCTCTTCTCGCACCGGGCCA<u>CAATCCACTGGGGTCGCT</u>ATGA – 3’ and was synthesised as an ssDNA oligo (Sigma-Aldrich). The synthesised IS2404 synthetic positive was resuspended in nuclease-free water and diluted to 0.001 pM, with 2 µL being used for extraction and positive controls. To confirm the presence of a true positive as opposed to contamination, 5 µL of the qPCR product was added to 1 µL of DNA Gel Loading Dye 6X (Thermo Scientific^TM^) and run on a 2 % agarose gel (TopVision Agarose Tablets, Thermo Scientific^TM^), with 1% SYBR Safe DNA Gel Stain (Invitrogen^TM^). The size of any positive IS2404 detection was assessed against 2 µL of 100 bp DNA Ladder (Promega) and run at 50 V for 1.5 hrs before being visualised with an EZ Gel Documentation System (Bio-Rad). Before the screening of insects commenced, the synthetic positive control for IS2404 was successfully designed and tested. By running the amplified PCR products on an agarose gel with the synthetic positive control and a real positive control, visual differentiation between a synthetic positive occurring at 120 bp and a true positive at 59 bp (Figure S1).

### Screening insects by qPCR for *M. ulcerans*

The qPCR screening was performed using three independent assays IS2404, IS2606 and KR ^16^. All samples were first screened with the IS2404 qPCR; if a positive was detected, additional confirmation was attempted with IS2606 and KR qPCR assays. Reactions were performed using 7.5 µL TaqMan^TM^ Fast Universal PCR Master Mix (2X), no AmpErase^TM^ UNG (Applied Biosystems^TM^), 1 µL of the primer-probe mix, 2 µL of DNA and 4.5 µL of nuclease-free water. A final primer-probe concentration for the IS2404 assay was as follows: 250:650:450nM for the forward primer, reverse primer and probe and 800:800:220nM for the forward primer, reverse primer and probe for the IS2606 and KR assay. A 2 µL volume of the synthetic positive control was added for the IS2404 reactions, whereas 2 µL of *M. ulcerans* DNA was used for IS2606 and KR. All reactions included six no-template extraction controls and were run in a 96 well plate format. Cycling conditions were as follows: denaturation at 95°C for 2 mins followed by 45 cycles at 95°C for 10 sec and 60°C for 30 sec, with qPCR performed on a QuantStudio^TM^ 5 Real-Time PCR System (Applied Biosystems^TM^). Positives were classified as reactions that produced a cycle quantification (Cq) value less than 40. Data were analysed using the QuantStudio^TM^ Design and Analysis Software v1.4.3 with the Delta Rn threshold set at 0.04 for IS2404 and IS2606 and Delta Rn threshold of 0.1 for KR. The maximum likelihood estimate (MLE) per 1,000 mosquitoes tested (bias corrected MLE for point estimation of infection rate and a skew-corrected score confidence intervals) was calculated from the pooled samples as described ^31^. Fisher’s Exact test for assessing the significance of differences in IS2404 PCR positivity between mosquito species was calculated in R 4.0.2 ^32^.

All qPCR screening was performed blind with mixed-species 96-well plates. The synthetic positive control was added at the DNA extraction phase to one well of each plate and to qPCR plates to check that both extraction and qPCR detections were successful. All no-template controls (extraction and qPCR stage) were checked to ensure they remained negative, and that synthetic positive controls were detected for both the DNA extraction and qPCR stage in each run. Positive samples were run on agarose gels to confirm they were true positives and not contamination from the synthetic positive control.

An IS2404 qPCR standard curve was prepared using 10-fold serial dilutions of *M. ulcerans* genomic DNA, with quadruplicate testing of each dilution. The DNA was extracted from *M. ulcerans* JKD8049 as described, and quantified using fluorimetry (Qubit, dsDNA HS ThermoFisher Scientific) ^22^. A limit-of-detection was defined as the lowest dilution that returned a positive signal for all four replicates. Genome equivalents were calculated based on the estimated mass of the *M. ulcerans* genome of 5.7 femtograms ^22^. IS*2404* cycle threshold (Ct) values were converted to genome equivalents (GE) to estimate bacterial load within a sample by reference to a standard curve (r^2^ = 0.9956, y = [-3.829Ln(x)+37.17]*Z, where y = Ct and x = amount of DNA [fg] and Z = the dilution factor]) (Figure S2). An IS2404 qPCR standard curve was fitted using non-linear regression in GraphPad Prism (v9.5.1).

### Mycobacterium ulcerans genome sequencing

Whole-genome sequencing was performed directly on DNA extracted from selected PCR positive mosquito samples and possum excreta specimens using a hybridisation capture approach, based on 120 nucleotide RNA baits spanning the 5.8 Mbp chromosome of the the *M. ulcerans* JKD8049 reference genome [Bioproject ID: PRJNA771185] (Datafile S1) (SureSelect Target enrichment system, Agilent, Santa Clara, CA, USA) and the Illumina *Nextera Flex for Enrichment with RNA Probes* protocol ^33^. Resulting sequence reads were submitted to NCBI GenBank and are available under Bioproject PRJNA943595 (Table S2).

### *Mycobacterium ulcerans* SNP calling, SNP imputation and phylogenetic analysis

To compare genomic variations between *M. ulcerans* clinical isolate genome sequences and sequence capture enrichment datasets, we mapped the sequence reads and called nucleotide variations using *Snippy* (v4.4.5) against a finished *M. ulcerans* reference chromosome, reconstructed from a Victorian clinical isolate (JKD8049; GenBank accession: NZ_CP085200.1) (https://github.com/tseemann/snippy). While standard parameters, including a minimum coverage of 10x were used for the clinical isolates and two possum sequence capture datasets, the mosquito sequence capture datasets had lower read coverage, necessitating the adjustment of parameters. Thus, the minimum coverage threshold was lowered to 1x to facilitate SNP calling for the mosquito sequence capture datasets. The resulting SNPs were combined with 117 SNPs obtained from a reference set of 36 *M. ulcerans* genomes that represented the previously defined population structure of the pathogen in Victoria ^35^. Due to the low read coverage however, the number of core variable nucleotide positions (VNPs) mapped among the five sequence capture datasets was variable (range: 22 −112 VNPs).

To enable inclusion of the sequence capture datasets that had missing SNP sites we employed a multivariate imputation approach, utilising the *IterativeImputer* function from *scikit-learn*^36^.

The combined alignment of 117 core genome SNPs from the sequence capture datasets and clinical isolate genomes served as the foundation for inferring a maximum likelihood phylogeny. This phylogeny was established using the GTR model of nucleotide substitution and executed with FastTree (v2.1.10) ^34^. The incorporation of R packages phytools (v1.0-1) ^37^and mapdata (v2.3.1) ^38^ allowed for the alignment of tree tips against a base map, facilitating the visualisation of geographical origins of the samples. Further details, including the code employed for missing SNP imputation and phylogeographic analysis, can be found in our GitHub repository: https://github.com/abuultjens/Mosquito_possum_human_genomic_analysis.

### *Aedes notoscriptus* typing and species confirmation sequencing

Mosquito genotyping was performed by sequence comparisons of a partial fragment of the cytochrome c oxidase subunit I (COI) gene ^30^. DNA was extracted using the above protocols. PCR was performed using 5 µL of 5x MyFi Reaction Buffer, 1 µL of MyFi DNA polymerase, 5 µL of DNA, and primer concentrations as described ^39^, with reaction made up to 25 µL with nuclease-free water. Reaction conditions were as follows for COI, initial denaturation at 95 °C for 1 min, followed by 35 cycles at 95°C for 20 sec, 46°C for 20 sec, and 72°C for 60 sec before a final extension at 72°C for 5 mins. A 5µL volume of the amplified PCR product was added to 1 µL of DNA Gel Loading Dye 6X (Thermo Scientific^TM^) and run on a 1 % agarose gel (TopVision Agarose Tablets, Thermo Scientific^TM^), with 1% SYBR Safe DNA Gel Stain (Invitrogen^TM^). 2 µL of 100 bp DNA Ladder (Promega) was added to confirm amplicons size and run at 100 V for 45 mins. PCR products that produced bands of the correct size were purified using the ISOLATE II PCR and Gel Kit (Bioline) as per manufacturer’s protocol and submitted for sequencing using an Applied Biosystems 3730xl capillary analyser (Macrogen), with sequencing occurring on both strands. Sequences were analysed in Geneious Prime (v2019.2.1) and trimmed to high-quality bases, aligned using ClustalW v2.1 and trimmed to a consensus region, *Ae. notoscriptus* COI (874 bp) and for species identification COI (816-882 bp). Sequences were analysed using blastn against the NCBI database. COI sequences generated for species identification are available under accession numbers OQ600123-4 and COI sequences for *Ae. notoscriptus* phylogenetics under accession numbers OQ588831-67 (Table S2).

### Mosquito bloodmeal analysis

Ninety blood-fed mosquitoes were identified as having an engorged abdomen and stilling having a red pigment (Sella score 2-3) indicating a fresh bloodmeal and were dissected with a sterile scalpel blade. Blood from the dissected abdomen was absorbed onto a 3 x 20 mm piece of a Whatman FTA^TM^ card (Merck) and placed in a 2 mL tube. DNA was extracted from the FTA^TM^ card using an ISOLATE II Genomic DNA Kit (Bioline) with a pre-lysis in 180 µL of Lysis Buffer GL and 25 µL Proteinase K for 2 hours before completing the extraction as per manufacturer’s protocol. Extracted DNA was amplified for Cyt *b* using primers previously described (Townzen et al. 2008), with MyTaq^TM^ HS Red Mix (Bioline), thermocycling conditions used were as follows: 95°C for 1 min, 30 cycles of 95°C for 15 secs, 50°C for 20 sec, 72°C for 20 sec, with a final extension at 72°C for 2 mins. Negative extraction and negative PCR controls were included with each PCR reaction. 5 µL of the amplified products was run on a 1% agarose gel containing 0.1% SYBR Safe DNA gel stain (Invitrogen) and visualised with a G:BOX Syngene blue light visualisation instrument. If a band was visualised at ∼480 bp the PCR product was purified using an ISOLATE II PCR and Gel Kit (Bioline).

PCR products were quantified using a Qubit (Invitrogen) Fluorometer with an HS dsDNA kit. Sequencing libraries were prepared using 10 ng of purified PCR product with a NEXTFLEX® Rapid XP DNA-Seq Kit (PerkinElmer) barcoded using NEXTFLEX® UDI Barcodes (PerkinElmer). As a result, 70 bloodmeal libraries were sequenced, as well as 6 PCR negative control and 3 extraction negatives on a NovaSeq 6000 (Illumina), with 2 Gb requested per sample.

Sequence data were analysed by identifying poor-quality reads using Rcorrector (Song & Florea, 2015) and removed with TrimGalore v0.6.5 (Krueger 2019). De novo assembly was performed on the remaining sequence reads using Trinity v2.8.6 (Grabherr et al., 2011). The resulting contigs were filtered to be between 400-480bp and analysed using BLASTN (Altschul et al. 1990) against the nucleotide database publicly available on the National Centre for Biotechnology Information (NCBI) website. The resulting hits were filtered to exclude those sequences which had < 97% sequence identity to the database. Contigs identified as a host bloodmeal were confirmed by mapping raw reads using BWA-MEM v0.7.17 (Li & Durbin, 2009). Sequence reads were submitted to Genbank (Table S2).

### Mosquito phylogenetic analysis

Mosquito genotyping was performed with COI as this genetic marker has previously been used to identify potential cryptic species within *Ae. notoscriptus* and can provide better resolution than other markers such as ND5, CAD or EPIC (Exon-Primed Intro Crossing) markers ^39^. Phylogenetic analysis was performed on trimmed consensus regions of *Ae. notoscriptus* COI. The substitution model was selected using jModelTest2 v2.1.10, with the topology taking the best of nearest neighbour interchange, subtree pruning and regrafting ^40^. The most appropriate substitution model was selected based on the Akaike information criterion. Maximum-likelihood trees were constructed in PhyML v3.3.2 with 1,000 bootstrap replicates; the gamma distribution parameter was used to estimate rate variation across sites ^41^. The Hasegawa-Kishino-Yano (HKY) substitution model was selected for the COI tree.

### Geographical data acquisition and spatial cluster analysis

The population map was created in qGIS v2.18.20 ^42^, using a 1 km^2^ population grid ^43^ with 2011 Victorian mesh block data. Since 2004, Buruli ulcer has been a notifiable condition in Victoria, requiring Health Department reporting by doctors and laboratories. De-identified case notification data of Buruli ulcer patients who had laboratory-confirmed *M. ulcerans* infection and who lived on the Mornington Peninsula during the years 2019-2020 were provided by the Victorian Department of Health. The cases were defined as patients with a clinical lesion that was diagnosed using IS2404 qPCR and culture ^16^. To conduct high-resolution spatial analyses, the data were aggregated at the mesh block level, the smallest geographical census units which typically contain 30-60 dwellings. The 2011 Victorian mesh block boundaries and the Victorian mesh block census population counts datasets were obtained from the Australian Bureau of Statistics (ABS) website ^44^. The datasets were joined using the unique mesh block IDs using the QGIS v.3.16.7 geographic information system software ^45^. The latitude and longitude (projected in GDA94) were derived from the centroids of the mesh block polygon. The dataset was then down-sampled to include only the Mornington Peninsula study area, specifically the Point Nepean and Rosebud-McCrae ABS level 2 Statistical Area.

SaTScan version 10.1.0 ^46^ was utilised to identify spatial clusters among trapped mosquitoes positive for *M. ulcerans*, possum excreta positive for *M. ulcerans*, and human Buruli ulcer cases. The software searches for instances where the observed number of spatial incidences exceeds the expected number within a circular window of varying size across a defined study area. A log likelihood ratio statistic is calculated for each window by comparing the number of observed and expected cases inside and outside the circle against the assumption of randomly distributed cases. In addition to the most likely cluster, there are usually secondary clusters with almost as high likelihood that substantially overlap with the primary clusters. These secondary clusters can be indicative of sub-clusters within the primary cluster or potentially distinct clusters that are spatially adjacent to the primary cluster. The Mornington Peninsula surveillance data used in these analyses consisted of trapped mosquitoes (177 traps screened for IS2404 collected: 12/11/19 to 20/03/20), *M. ulcerans* detected in possum excreta collected during the summer (December to February) of 2019 using data from a previous study ^47^ and notified human Buruli ulcer cases from the study area in the years 2019-2020. The use of possum excreta collected during the summer was appropriate as Buruli ulcer transmission is most likely to occur during that time of year ^48,49^. For each of the three data sources (trapped mosquitoes, possum excreta, and human cases), the null hypothesis assumes that *M. ulcerans* detections or Buruli ulcer cases are uniformly distributed across the study area, where the alternative hypothesis suggests that there may be certain locations with higher rates than expected if the risk was evenly distributed. Primary and secondary clusters were accepted only if the secondary clusters did not overlap with previously reported clusters with a higher likelihood. Given that the trapped mosquito and possum excreta IS2404 PCR results were binary (positive or negative), a Bernoulli model was used to scan for spatial clusters, with the maximum cluster size set to 50% of the population size. The human Buruli ulcer case data aggregated at the mesh block level varied in number, with some mesh blocks having zero cases and others having one or more. We applied the Poisson probability model to the notified Buruli ulcer case counts, using a background population at risk that was derived from the 2011 population census. The maximum cluster size was limited to 14,481 individuals, 10% of the total population at risk. To determine the likelihood of a triple-cluster overlap between the three SaTScan analyses occurring by chance, we conducted a permutation test (https://github.com/abuultjens/BU-3-way-SatScan). In each of 10,000 iterations, the geographical coordinates for each variable were randomly shuffled. The number of SaTScan clusters with triple overlap was determined using the ‘sf’ package ^50^ in the R statistical programming language ^51^.

## Results

### Mosquito surveys of the Mornington Peninsula

A primary goal of this research was to test the hypothesis that mosquitoes are associated with *M. ulcerans* transmission, as reflected by IS2404 PCR positivity at a certain frequency in areas of the Mornington Peninsula with human Buruli ulcer cases. A total of 73,580 mosquitoes were collected, consisting of 72,263 females (Table 2) and 1,317 males (Table S2). The majority (90%) of these insects were collected in the large survey of 2019-2020 (Datafile S2). Across all five surveys, 26 different mosquito species were collected covering six genera (Table 2). The most dominant species identified during the 2019-2020 survey was *Cx. molestus* accounting for 42% of mosquitoes, followed by *Ae. notoscriptus* (35%) and *Cx. australicus* (8%). Twenty-three other species comprised the remaining 15% of mosquitoes. The distribution of the two dominant species across the survey area is shown in Figure 2. This mapping revealed an asymmetric distribution of each species, with *Ae. notoscriptus* dominant to the eastern end and *Cx. molestus* dominant towards the western end of the Peninsula (Figure 2).

**Figure 2.**
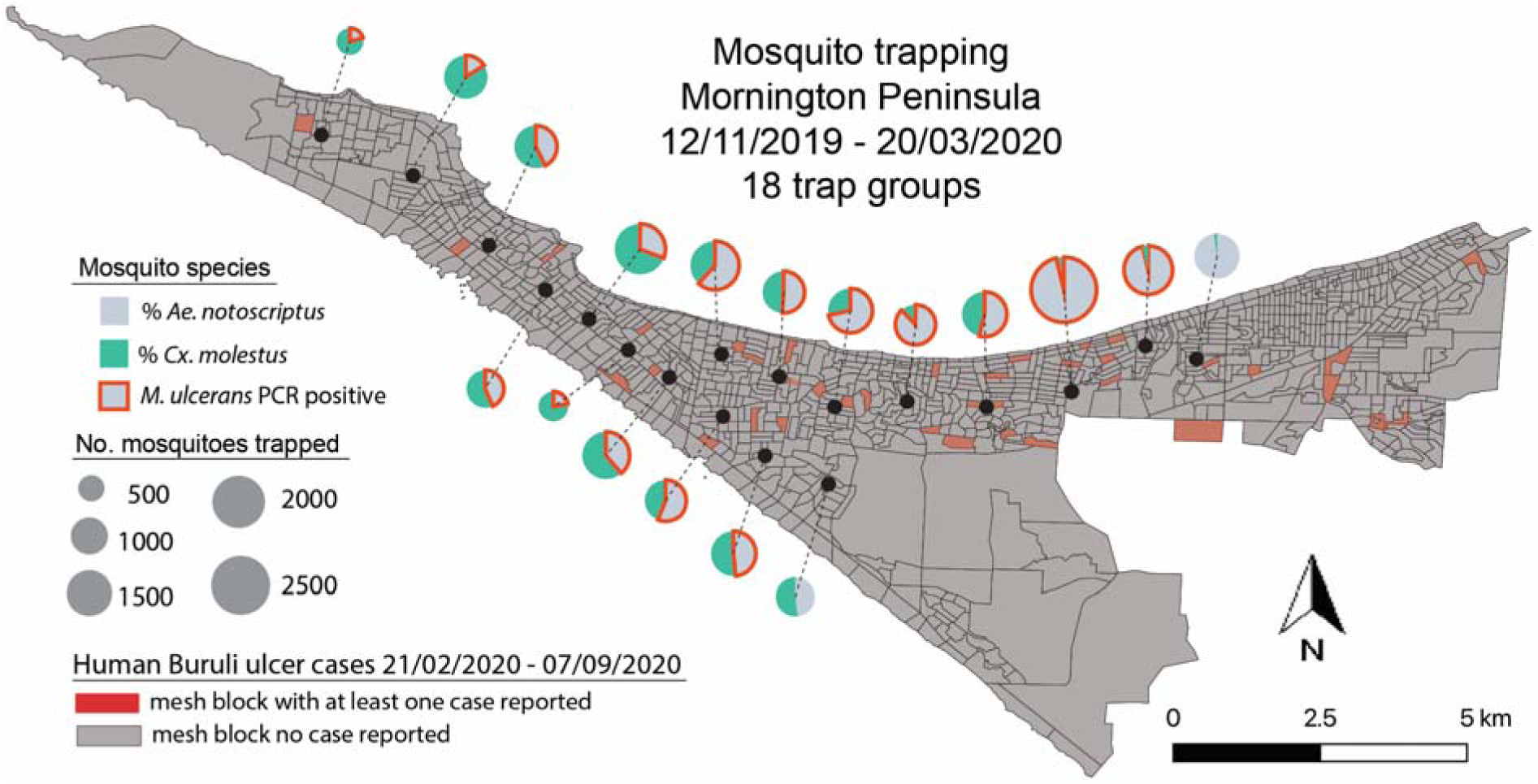
Dominant mosquito species distribution across the Mornington Peninsula. Map showing the proportional distribution of the two dominant mosquito species trapped during 2019/2020. The pie charts are an aggregation of the 180 different trap sites. Trap groups containing mosquitoes that were PCR positive for M. ulcerans are also indicated. Shown too, are the meshblock statistical areas, with those in red containing at least one human Buruli ulcer case diagnosed in 2019-2020.

**Table 2.**
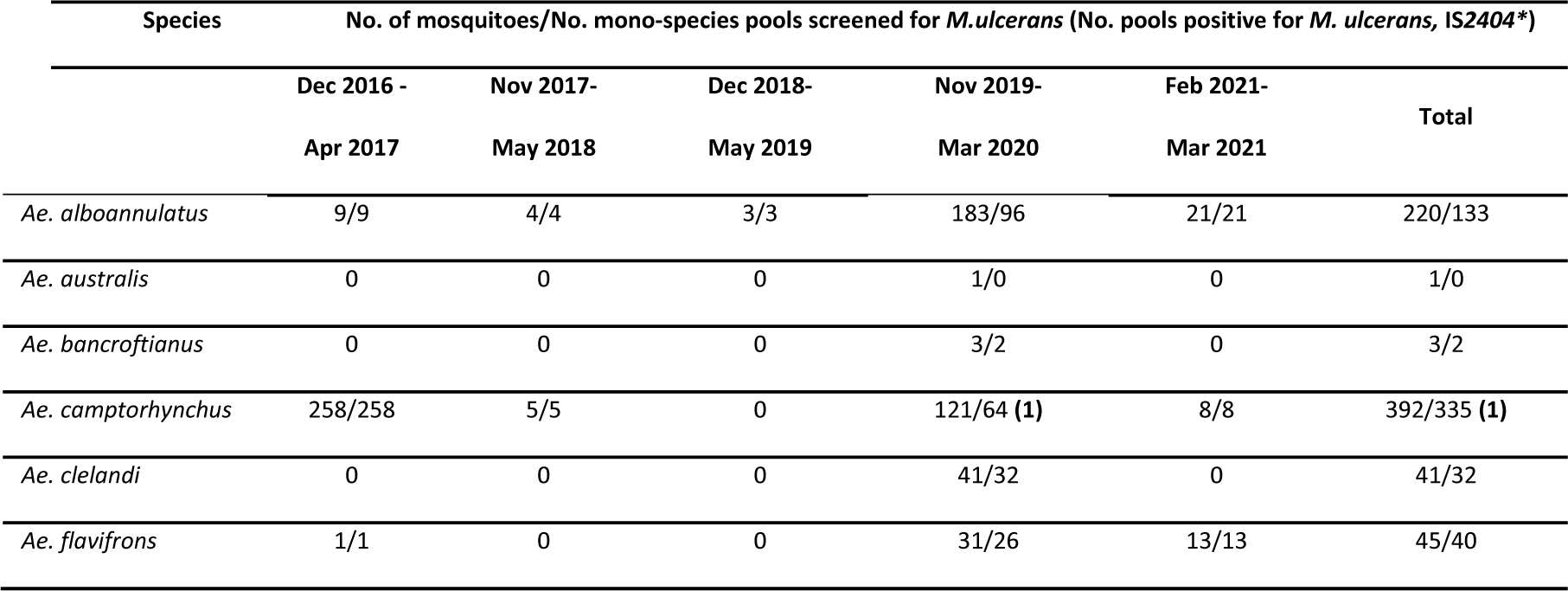
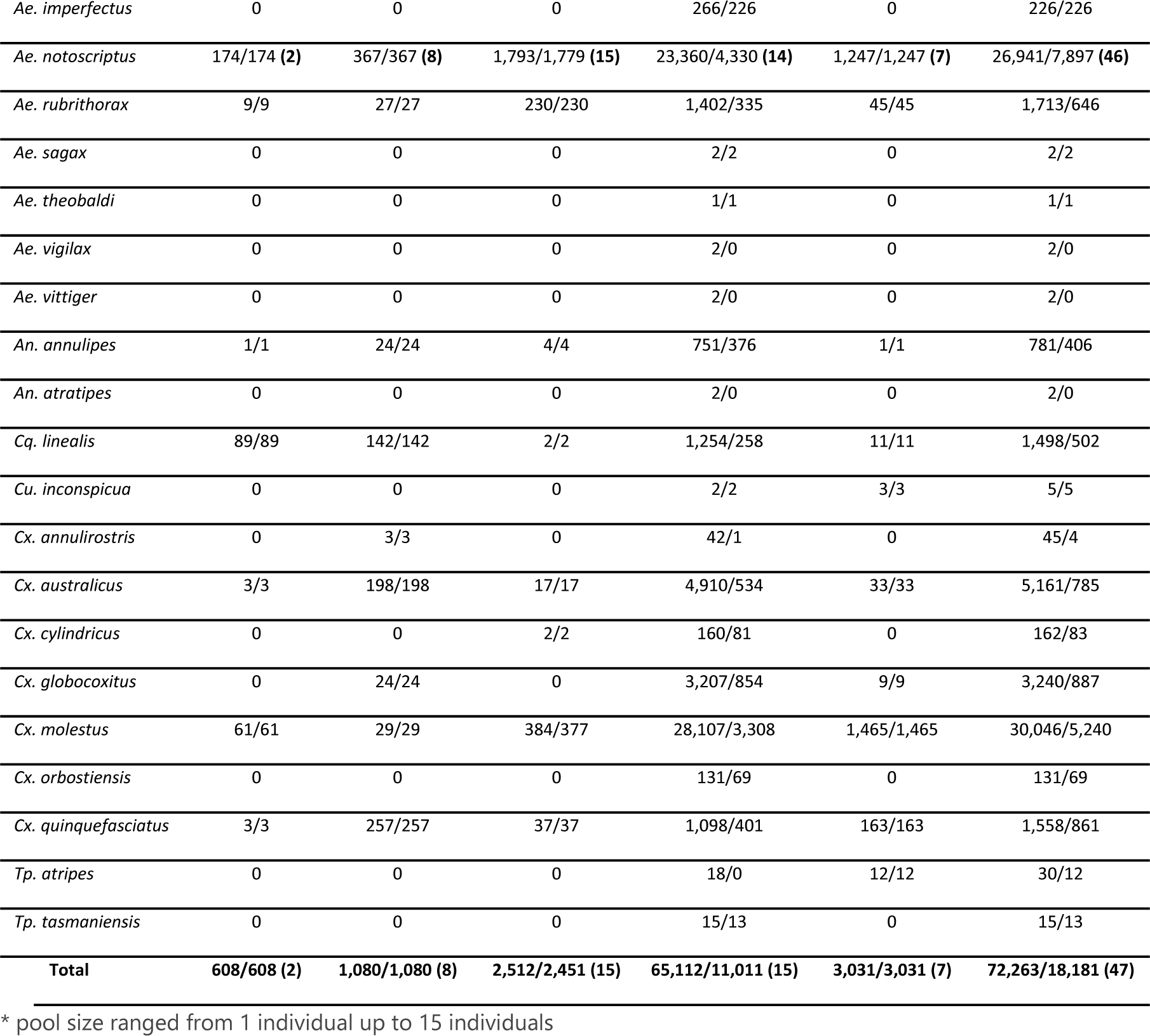
Female mosquitoes trapped on the Mornington Peninsula and screened by IS2404 PCR.

| Species | No. of mosquitoes/No. mono-species pools screened for *M. ulcerans* (No. pools positive for *M. ulcerans*, IS2404*) |  |  |  |  | Total |
| --- | --- | --- | --- | --- | --- | --- |
|  | Dec 2016 -<br>Apr 2017 | Nov 2017-May 2018 | Dec 2018-May 2019 | Nov 2019-Mar 2020 | Feb 2021-Mar 2021 |  |
| *Ae. alboannulatus* | 9/9 | 4/4 | 3/3 | 183/96 | 21/21 | 220/133 |
| *Ae. australis* | 0 | 0 | 0 | 1/0 | 0 | 1/0 |
| *Ae. bancroftianus* | 0 | 0 | 0 | 3/2 | 0 | 3/2 |
| *Ae. camptorhynchus* | 258/258 | 5/5 | 0 | 121/64 (1) | 8/8 | 392/335 (1) |
| *Ae. clelandi* | 0 | 0 | 0 | 41/32 | 0 | 41/32 |
| *Ae. flavifrons* | 1/1 | 0 | 0 | 31/26 | 13/13 | 45/40 |
| *Ae. imperfectus* | 0 | 0 | 0 | 266/226 | 0 | 226/226 |
| *Ae. notoscriptus* | 174/174 (2) | 367/367 (8) | 1,793/1,779 (15) | 23,360/4,330 (14) | 1,247/1,247 (7) | 26,941/7,897 (46) |
| *Ae. rubrithorax* | 9/9 | 27/27 | 230/230 | 1,402/335 | 45/45 | 1,713/646 |
| *Ae. sagax* | 0 | 0 | 0 | 2/2 | 0 | 2/2 |
| *Ae. theobaldi* | 0 | 0 | 0 | 1/1 | 0 | 1/1 |
| *Ae. vigilax* | 0 | 0 | 0 | 2/0 | 0 | 2/0 |
| *Ae. vittiger* | 0 | 0 | 0 | 2/0 | 0 | 2/0 |
| *An. annulipes* | 1/1 | 24/24 | 4/4 | 751/376 | 1/1 | 781/406 |
| *An. atratipes* | 0 | 0 | 0 | 2/0 | 0 | 2/0 |
| *Cq. linealis* | 89/89 | 142/142 | 2/2 | 1,254/258 | 11/11 | 1,498/502 |
| *Cu. inconspicua* | 0 | 0 | 0 | 2/2 | 3/3 | 5/5 |
| *Cx. annulirostris* | 0 | 3/3 | 0 | 42/1 | 0 | 45/4 |
| *Cx. australicus* | 3/3 | 198/198 | 17/17 | 4,910/534 | 33/33 | 5,161/785 |
| *Cx. cylindricus* | 0 | 0 | 2/2 | 160/81 | 0 | 162/83 |
| *Cx. globocoxitus* | 0 | 24/24 | 0 | 3,207/854 | 9/9 | 3,240/887 |
| *Cx. molestus* | 61/61 | 29/29 | 384/377 | 28,107/3,308 | 1,465/1,465 | 30,046/5,240 |
| *Cx. orbostiensis* | 0 | 0 | 0 | 131/69 | 0 | 131/69 |
| *Cx. quinquefasciatus* | 3/3 | 257/257 | 37/37 | 1,098/401 | 163/163 | 1,558/861 |
| *Tp. atripes* | 0 | 0 | 0 | 18/0 | 12/12 | 30/12 |
| *Tp. tasmaniensis* | 0 | 0 | 0 | 15/13 | 0 | 15/13 |
| **Total** | **608/608 (2)** | **1,080/1,080 (8)** | **2,512/2,451 (15)** | **65,112/11,011 (15)** | **3,031/3,031 (7)** | **72,263/18,181 (47)** |

\* pool size ranged from 1 individual up to 15 individuals

### M. ulcerans PCR positive mosquitoes were predominantly Aedes notoscriptus

The IS2404 qPCR assay was used to infer the presence of *M. ulcerans* in association with mosquitoes ^16^. Of the 73,580 mosquitoes trapped across all years, 18,610 (25%) were screened by IS2404 qPCR for the presence of *M. ulcerans* (Table 2). *M. ulcerans* qPCR positives were observed in 53 pools or individuals of *Ae. notoscriptus*, with detections occurring in each year across all survey events (Figure 2, Table 2). Only one other mosquito species tested IS*2404* positive, which was a two-insect pool of *Ae. camptorhynchus* (Table 3). The positive association between *M. ulcerans* and *Ae. notoscriptus* compared to other mosquitoes, in particular the other abundant mosquito species, *Cx. molestus*, was highly significant (p<0.0001, Fisher’s exact test).

**Table 3.**
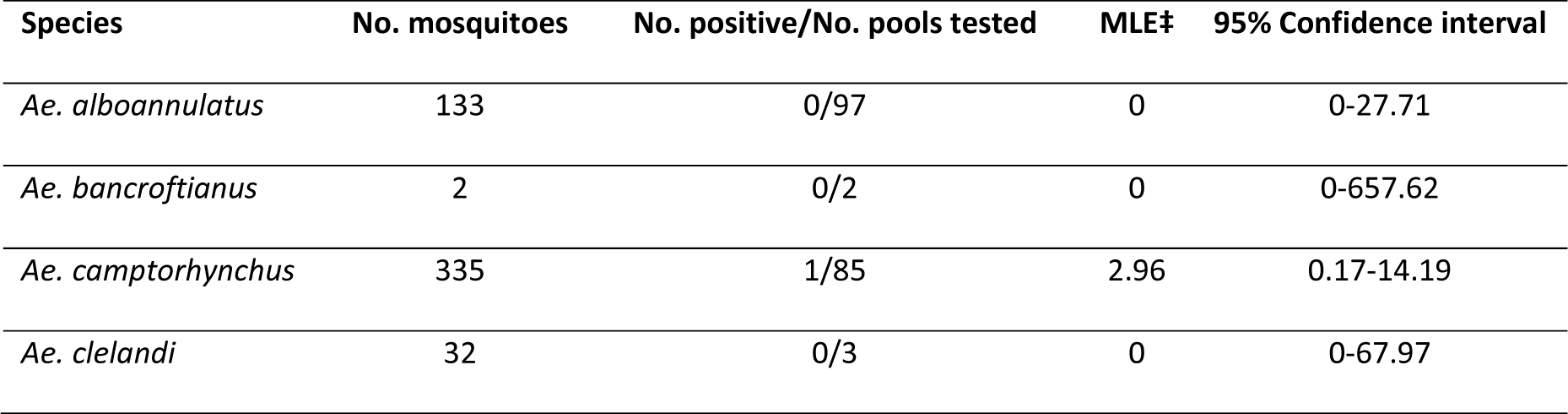
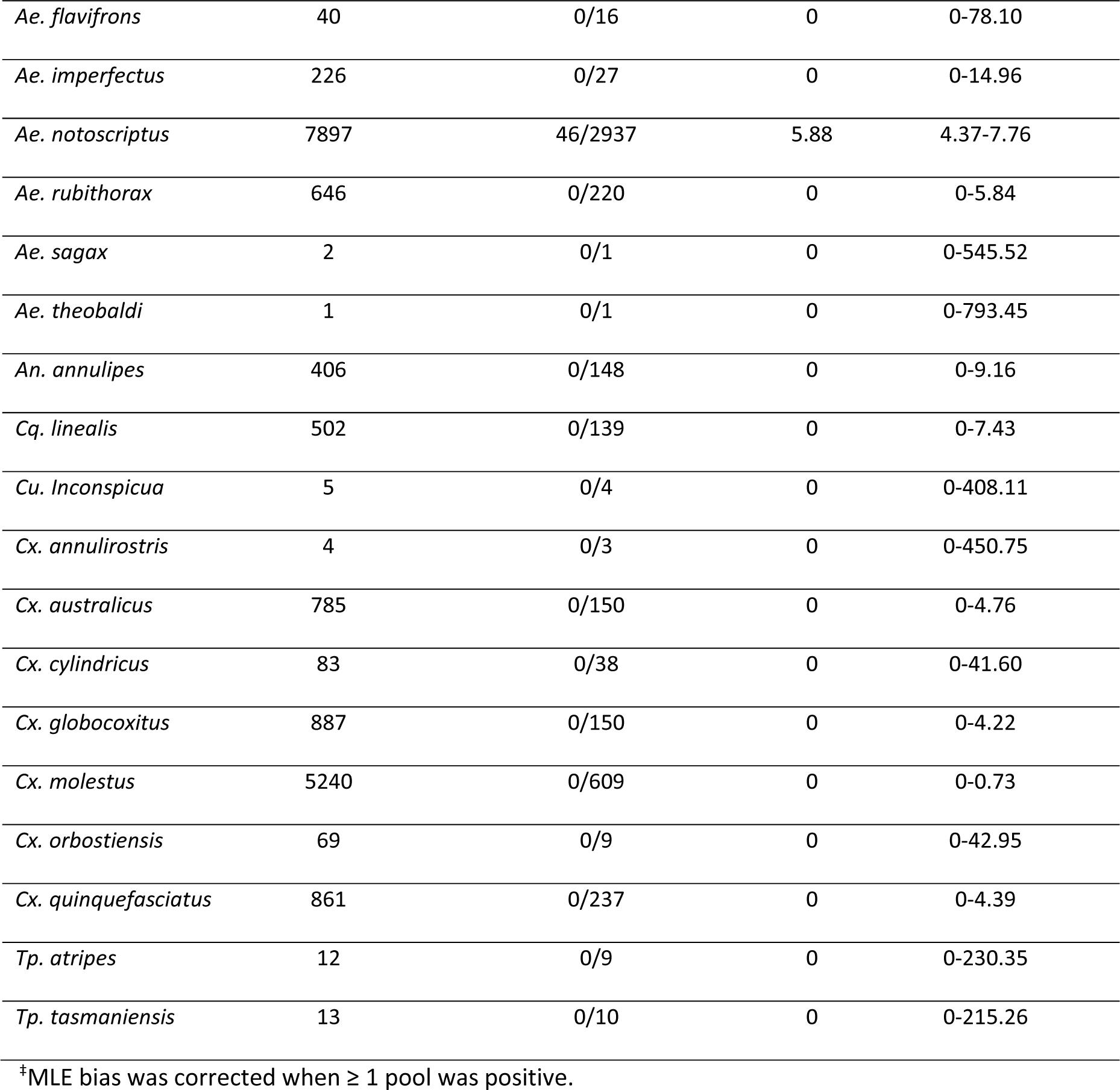
The number *of M. ulcerans* positive pools tested by insertion sequence IS2404 qPCR and Maximum likelihood estimate (MLE) of *M. ulcerans* per 1,000 mosquitoes trapped in the Mornington.

| Species | No. mosquitoes | No. positive/No. pools tested | MLE‡ | 95% Confidence interval |
| --- | --- | --- | --- | --- |
| *Ae. alboannulatus* | 133 | 0/97 | 0 | 0-27.71 |
| *Ae. bancroftianus* | 2 | 0/2 | 0 | 0-657.62 |
| *Ae. camptorhynchus* | 335 | 1/85 | 2.96 | 0.17-14.19 |
| *Ae. clelandi* | 32 | 0/3 | 0 | 0-67.97 |
| *Ae. flavifrons* | 40 | 0/16 | 0 | 0-78.10 |
| *Ae. imperfectus* | 226 | 0/27 | 0 | 0-14.96 |
| *Ae. notoscriptus* | 7897 | 46/2937 | 5.88 | 4.37-7.76 |
| *Ae. rubithorax* | 646 | 0/220 | 0 | 0-5.84 |
| *Ae. sagax* | 2 | 0/1 | 0 | 0-545.52 |
| *Ae. theobaldi* | 1 | 0/1 | 0 | 0-793.45 |
| *An. annulipes* | 406 | 0/148 | 0 | 0-9.16 |
| *Cq. linealis* | 502 | 0/139 | 0 | 0-7.43 |
| *Cu. inconspicua* | 5 | 0/4 | 0 | 0-408.11 |
| *Cx. annulirostris* | 4 | 0/3 | 0 | 0-450.75 |
| *Cx. australicus* | 785 | 0/150 | 0 | 0-4.76 |
| *Cx. cylindricus* | 83 | 0/38 | 0 | 0-41.60 |
| *Cx. globocoxitus* | 887 | 0/150 | 0 | 0-4.22 |
| *Cx. molestus* | 5240 | 0/609 | 0 | 0-0.73 |
| *Cx. orbostiensis* | 69 | 0/9 | 0 | 0-42.95 |
| *Cx. quinquefasciatus* | 861 | 0/237 | 0 | 0-4.39 |
| *Tp. atripes* | 12 | 0/9 | 0 | 0-230.35 |
| *Tp. tasmaniensis* | 13 | 0/10 | 0 | 0-215.26 |

<sup>‡</sup>MLE bias was corrected when $\geq 1$ pool was positive.

Subsets of *Ae. notoscriptus* were screened individually to better estimate the prevalence of *M. ulcerans* positive mosquitoes in this species. Individually tested mosquitoes included all of the 2017/2018 collection comprising 367/367 individuals with 8 positives (2.2%), 448/4,330 of the 2019/2020 collection with 3 positives (0.67%), and all 1,247 individuals of the 2021 collection with 7 positives (0.56%) (Table S3). Thus, based on screening individual insects, 18/2,062 (0.87%) *Ae. notoscriptus* were IS2404 PCR positive. All other *Ae. notoscriptus* and other mosquito species in this study were screened in pools (up to 15 insects per pool) with 24/747 *Ae. notoscriptus* pools (3%), PCR positive (Table S4).

Of the 46 *Ae. notoscriptus* that tested positive for IS2404, 26 (56%) were confirmed positive for IS2606 and 31 (67%) for KR, with 24 (52%) pools positive by all three qPCR assays (Table S3). Of the 46 IS2404 positive *Ae. notoscriptus*, eight (17%) pools were not tested using the IS2606 assay and one (2%) pool using the KR assay due to limited template DNA available. The average Ct value for IS2404 positive *Ae. notoscriptus* was 36.00 (range 29.74-39.65). The average Ct value for IS2606 was 37.73 (range 32.49-45.00), and the average Ct value for KR was 34.19 (range 28.52-39.26). With reference to an IS2404 qPCR standard curve (Figure S2), we estimated the *M. ulcerans* burden per mosquito. The mean bacterial genome equivalents (GE) per insect was 294 GE (range: 11 – 4200) (Table S3). The MLE of estimated infection rate for all *Ae. notoscriptus* was 5.88 based on the IS2404 *M. ulcerans* qPCR-positive mosquitoes per 1,000 tested (95% CI 4.37-7.76) for all *Ae. notoscriptus* tested over the years and 2.96 for *Ae. camptorhynchus* (95% CI 0.17-14.19) (Table 3).

### IS2404 PCR positive Ae. notoscriptus and possum excreta yield M. ulcerans genomic SNP profiles that match human patient isolates

Genomic epidemiological studies have shown there are characteristic SNP signatures associated with *M. ulcerans* clinical isolates from specific endemic areas in southeastern Australia ^35^. To test if the *M. ulcerans* genotypes present in mosquitoes from our study area matched that found in possum excreta and human Buruli ulcer cases in the same region, we conducted genome sequence enrichment. This method facilitates whole genome sequence analysis of target pathogen genomes directly from complex microbial samples, when competing background DNA precludes sequencing the target pathogen genome directly. As reported in other sequence enrichment studies, we observed decreasing genome sequence recovery with decreasing pathogen load, as indicated by increasing IS2404 Ct values (Figure 3A) ^52^. Nevertheless, DNA sequence reads were obtained from five IS2404 positive mosquitoes and two IS2404 positive possum excreta specimens, the latter specimens collected as part of a large field survey of *M. ulcerans* in Australian native possum excreta in the region ^53^. While DNA sequence reads were obtained across the length of the 5.6Mbp *M. ulcerans* reference chromosome, for all five mosquitoes and the two possum excreta specimens, only the excreta specimens (IS2404 Ct values <23) and three of the five mosquito DNA extracts (ID: 5675, Ct 32.62; ID: 226, Ct 33.47; ID: 819, Ct 31.20) yielded sufficient *M. ulcerans* reads for SNP-calling (Figure 3B).

**Figure 3.**
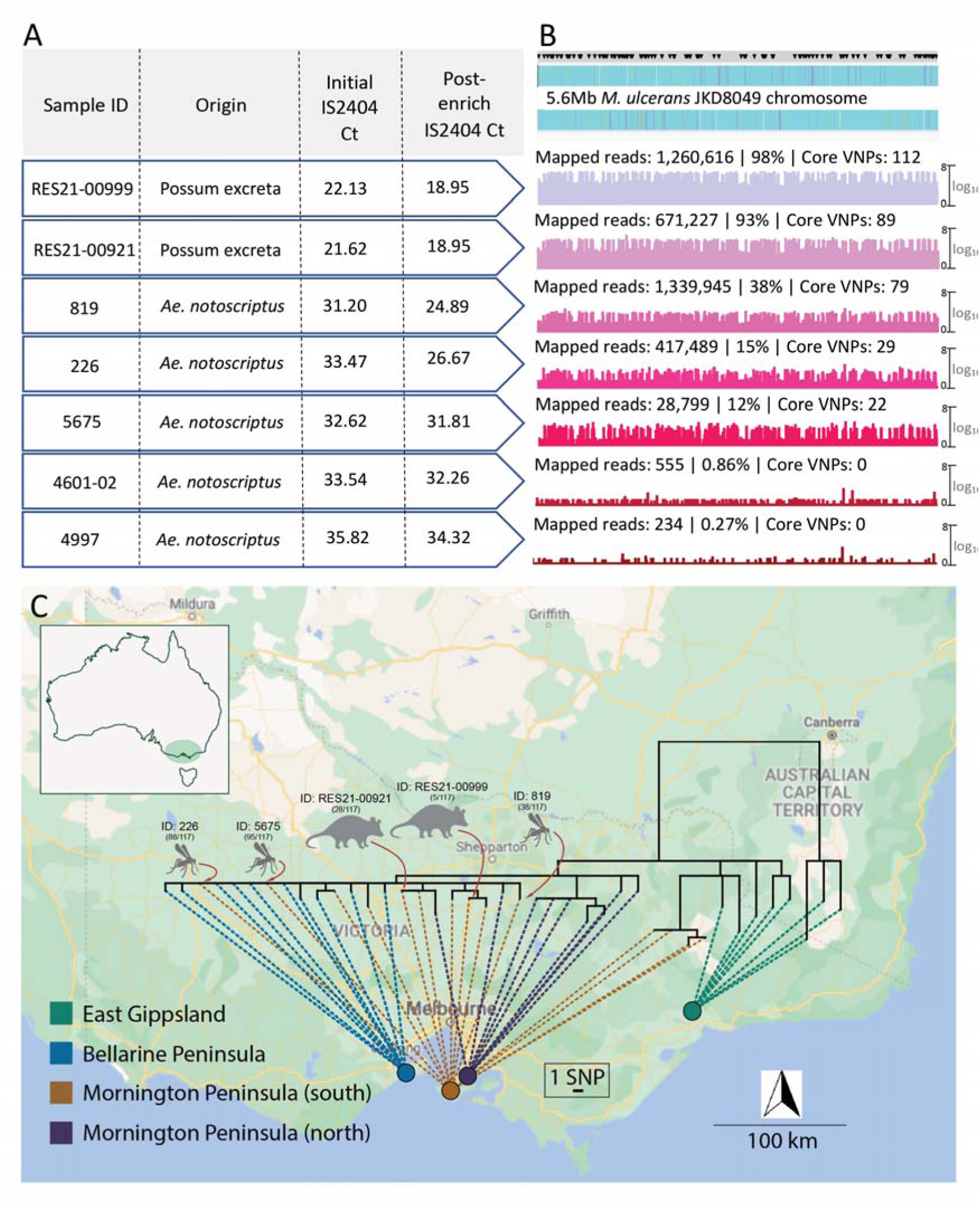
Comparison of M. ulcerans genome from human Buruli ulcer cases compared with sequences recovered from possum excreta and mosquitoes on the Mornington Peninsula. (A) Summary of IS2404 qPCR screening of primary samples and the sequence capture libraries pre- and post-enrichment. Note that possum excreta were enriched as barcoded sequence library pools so share the same pre- and post-enrichment Ct values; (B) Artemis coverage plots depicting sequence capture reads mapped to the M. ulcerans JKD8409 chromosome from possum excreta samples and three qPCR positive mosquitoes (labels are: numbers of mapped reads|percentage of total chromosome bases mapped|number of core Variable Nucleotide Positions [VNPs] covered). Grey horizontal bar above the above the chromosome map depicts the sites of 117 core SNPs (black inverted triangles); (C) Maximum-likelihood phylogeny inferred from an alignment of 117 core-genome SNPs using reads mapped to the JKD8049 reference chromosome, and with tips aligned with environmental sample origin or patient origin. Dataset includes reads from the five environmental samples listed in panel with >21 VNPs (A), and a reference collection of 36 M. ulcerans genomes representing the genomic diversity of the M. ulcerans population in southeastern Australia ^35^. The shortest vertical branch length represents a single SNP difference, as per the scale bar.

To assess the relatedness of *M. ulcerans* genomes from mosquito and excreta to human *M. ulcerans* isolates, we inferred a phylogeny using a 5.6Mbp local *M. ulcerans* reference chromosome and sequence reads from 36 published clinical *M. ulcerans* genomes from southeastern Australia ^35^, based on a core genome alignment with 117 SNP positions. These 36 genomes were selected because they spanned our previously reported population structure of *M. ulcerans* in this locale (Table S1) ^35^. Alignment of the five mosquito and two excreta sequence capture datasets revealed reads mapping across the length of the *M.ulcerans* chromosome - confirming the presence of the pathogen on (or in) *Ae. notoscriptus* and in the possum faecal material (Figure 3B).

Despite chromosome-wide read mapping for the seven sequence capture datasets, the relatively low and incomplete coverage meant that these datasets didn’t have reads spanning all 117 variable loci (range: 22-112 sites) (Figure 3B). Thus, to enable the inclusion of the sequence capture datasets into a phylogenomic analysis of 117 core SNPs, a multivariate statistical approach using an imputation model was employed (Figure S3A). The model was trained on the 117 SNPs from the core genome alignment of all 36 clinical isolates to predict with high confidence (impute) the missing SNP alleles of the sequence capture datasets (see methods). To validate the model, we initially masked 95 random SNP sites as missing data for a random set of five of the 36 clinical isolates, leaving 22 SNPs. Using the multivariate imputation model trained on the full 117 sites of the remaining 31 clinical isolate genomes, we predicted the missing alleles of the five masked clinical isolates (Figure S3A). This approach yielded a mean accuracy of 97% in correctly predicting missing alleles, given the 22 available SNPs. To ensure the model was robust, we randomly selected five *M. ulcerans* clinical isolate genomes and then varied the number of masked chromosome SNP sites to simulate missing data and replicated this process 100x. This analysis showed the high performance of the imputation model, with a mean accuracy of 94% when 100/117 SNP sites were randomly masked and then imputed (Figure S3B). Using the validated imputation model to predict the missing alleles for the five sequence capture datasets and create a sequence alignment (Dataset S3), a maximum likelihood tree was inferred using all 117 variable sites with location of tree tips aligned with geographic origin. This phylogeny showed possum excreta and mosquito *M. ulcerans* genotypes co-clustered with each other and with human *M. ulcerans* isolates from the Mornington Peninsula, with 0-5 SNP differences between any pairwise comparison (Figure 3C). This pattern is consistent with a shared transmission cycle between possums, mosquitoes and humans. Individual phylogenies inferred with and without SNP allele imputation showed that imputation did not create artifactual tree topologies (Figure S4).

### Genetic characterisation of Ae. notoscriptus collected on the Mornington Peninsula

To assess if *M. ulcerans* presence was associated with a particular clade of *Ae. notoscriptus* on the Mornington Peninsula we compared COI gene sequences for 18 *M. ulcerans* positive (confirmed by all IS2404, IS2606 and KR qPCR assays) *Ae. notoscriptus* and 19 *M. ulcerans* negative *Ae. notoscriptus* (Figure 2). Based on the COI phylogenetic tree, *Ae. notoscriptus* from the Mornington Peninsula spanned the three previously identified clades and there was no association between *M. ulcerans* and a particular mosquito lineage (Figure S5) ^39^.

### IS2404 PCR screening of arthropods other than mosquitoes

We also investigated the association of *M. ulcerans* with arthropods other than mosquitoes in the region using yellow sticky traps (YST) and sticky ovitraps (SO). A total of 21,000 specimens were collected and sorted from 278 YST and 33 SO traps. We were able to classify 2,696 as insects that may bite or pierce human skin (Table 4). Flies were the largest group collected on the sticky traps. YSTs collected more insects than the SOs, but this was proportional to the number of traps set (Table 4). Of the 2,696 insects screened by PCR, only two flies tested positive for *M. ulcerans* (Table S3). Both flies had high Ct values for IS2404 (Ct 35.92 and 37.54) and each sample was only confirmed for either IS2606 or KR assay (Table S3), but not with all three assays. Both insects were blowflies (*Calliphora hilli* Patton) based on morphology and sequencing of the COI region with sequences having >99.86 nt identity to *C. hilli* (Table S1).

**Table 4.** Number of arthropods collected on Yellow Sticky Traps (YST) and Sticky Ovitraps (SO) on the Mornington Peninsula that might bite or pierce human skin.

| Trap | Ants | Biting midges | Flies | Mosquitoes | Spiders | Ticks | Wasps | Total |
| --- | --- | --- | --- | --- | --- | --- | --- | --- |
| YST | 28 | 58 | 1,562 | 10 | 76 | 21 | 562 | 2,317 |
| SO | 70 | 1 | 144 | 11 | 141 | 4 | 8 | 379 |
| **Total** | **98** | **59** | **1,706** | **21** | **217** | **25** | **570** | **2,696** |

### Mosquito bloodmeal analysis

Of the mosquitoes collected, a proportion of individuals identified as being engorged (bloodfed) were PCR screened and the resulting amplicon sequenced for Cytochrome B (CytB) to identify host bloodmeal sources. A total of 90 individual engorged mosquitoes were extracted, with 70 DNA preparations producing high quality amplicons that were of sufficient concentration to permit Illumina amplicon sequencing. After quality filtering, 36 individuals were identified as having had a recent bloodmeal: 14 *Cx. molestus*, 13 *Ae. notoscriptus*, 2 *Ae. rubrithorax*, 2 *Cx. globocoxitus*, 2 *Cx. quinquefasciatus*, 2 *Cq. linealis* and 1 *Cx. australicus*. Of the bloodmeals detected, common ringtail possum was the most commonly identified with 20 detections across the 36 samples, followed by 17 blackbirds, 13 humans, 11 red wattle birds, and 5 little wattle birds; a further 16 bloodmeals identified 10 minor host species (Figure 4, Figure S6). Dual bloodmeals were commonly identified, with 55% (20/36) of individuals having more than one bloodmeal identified. Additionally, two mosquitoes had evidence of three different bloodmeal sources within an individual insect. Three individuals (2 x *Ae. notoscriptus* and 1 x *Ae. rubrithorax*) were also identified as having dual bloodmeals from ringtail possum and human origins (Figure 4).

**Figure 4.**
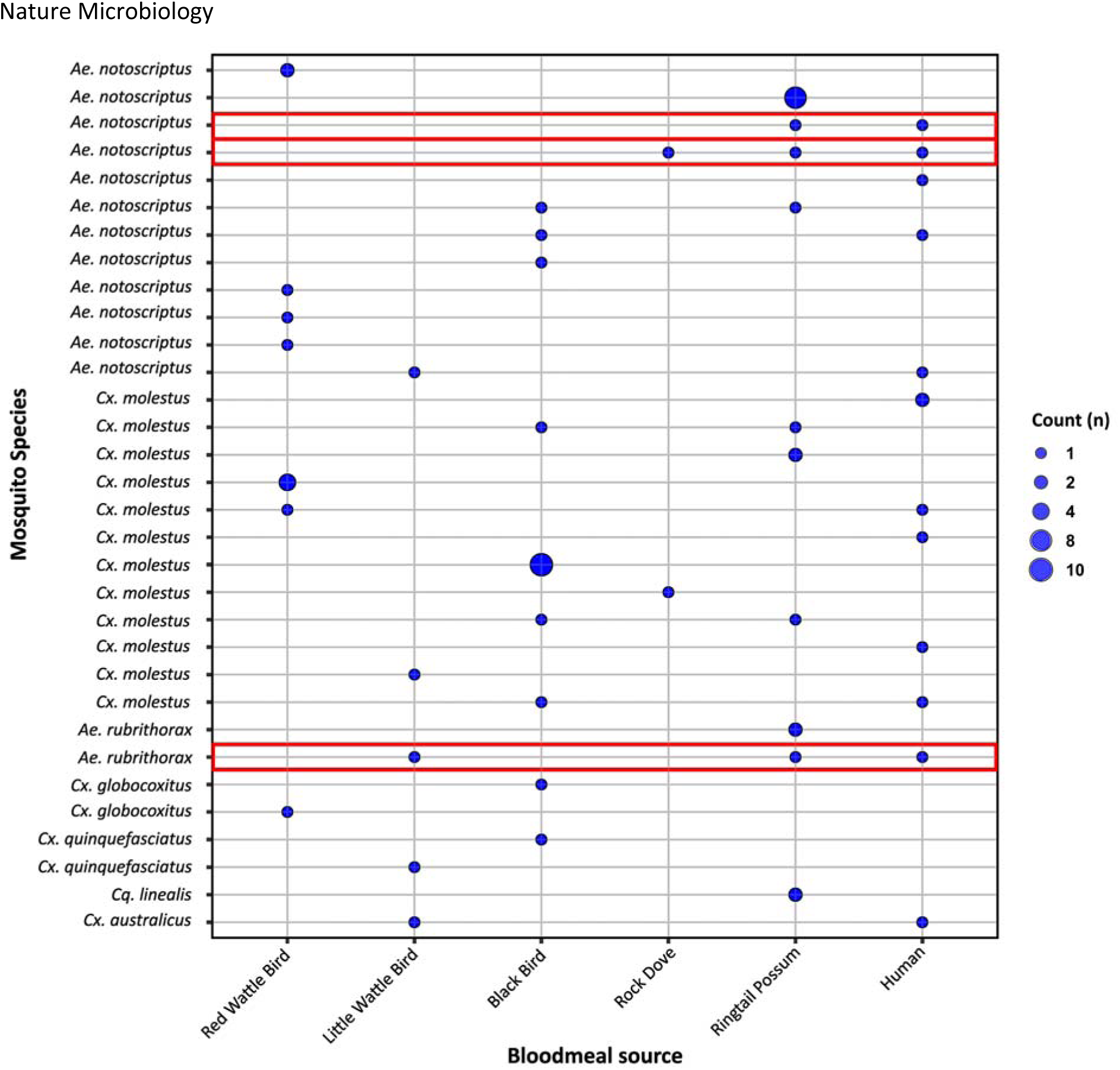
Mosquito blood meal analysis. Summary of cytochrome B (Cytb) gene sequences from the 36 bloodfed mosquitoes to identify host bloodmeal sources. A blue circle indicates positive for host blood source; the larger the circle, the more individual mosquitoes with an identical bloodmeal profile. Red boxes indicate individual mosquitoes which have dual bloodmeals for both humans and ringtail possum sources.

### Spatial clustering links *M. ulcerans* positive mosquitoes with positive possum excreta and human Buruli ulcer cases

In our previous research, we demonstrated a spatial association between possum excreta containing *M. ulcerans*, as detected by structured surveys, and clusters of human Buruli ulcer cases ^53^. To expand on this finding, we investigated whether further spatial clustering associations could be detected by analysing qualitative qPCR (IS2404) data of trapped mosquitoes using SaTScan. Here, three separate analyses were conducted: (i) human Buruli ulcer case data from individuals reported to have acquired the disease in the study area during 2019-2020, (ii) qPCR data from possum excreta collected in a previous investigation (2018-2019), and (iii) qPCR data from trapped mosquitoes (2019-2020). These analyses identified a single mosquito cluster, four possum clusters and six human Buruli ulcer clusters (Figure 5A-B). Notably, one human Buruli ulcer cluster and two possum excreta clusters had higher numbers of observed Buruli ulcer cases or *M. ulcerans* detections, respectively, than expected if uniformly distributed (p-value <0.05) (Figure 5B). Importantly, the analyses revealed an instance of triple cluster overlap (mosquito/possum excreta/human) in the Mornington Peninsula suburb of Rye, where all three SaTScan analyses had an overlapping cluster (Figure 5A).

**Figure 5.**
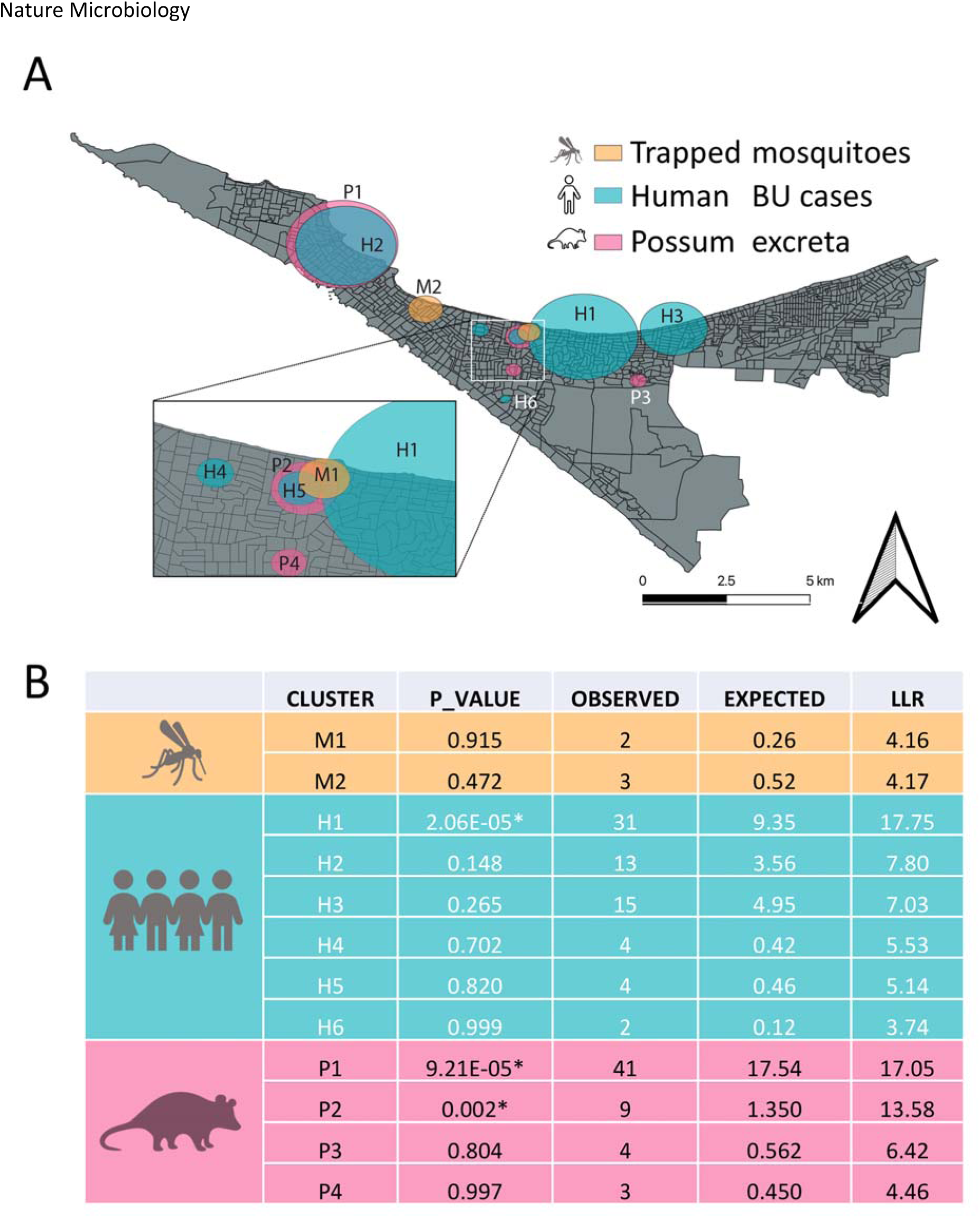
SaTScan analyses of trapped mosquitoes, possum excreta, and human Buruli ulcer cases on the Mornington Peninsula, revealing spatial clustering. A) Map illustrating the clusters identified by the three separate SaTScan analyses: (I) trapped mosquitoes (177 traps screened for IS2404 collected: 12.11.19 to 20.03.20), (ii) M. ulcerans detected in possum excreta collected during the summer of 2019 (Dec 2018 – Feb 2019) using data from a previous study, and (iii) notified human Buruli ulcer cases from the study area in the years 2019-2020. The zoomed insert highlights an instance where all three analyses had overlapping clusters in the suburb of Rye. B) Table summarising the SaTScan results for all identified clusters. Log likelihood ratio is abbreviated as “LLR.” Clusters with p-values <0.05 are marked with asterisks.

The results of our permutation test to explore the probability of these three categories (mosquitoes, possums and humans) overlapping showed that a triple-cluster overlap occurred with randomly rearranged location labels at a low frequency of 8.9%. This analysis adds support to a causal relationship between the presence of *M. ulcerans* in possums and mosquitoes and humans contracting Buruli ulcer in the Mornington Peninsula suburb of Rye.

## Discussion

Using insect field surveys augmented with genomics we have addressed *Barnett Criteria* 1 – 3 (Table 1) to add to a hierarchy of evidence indicting mosquitoes as principal vectors in the spread of *M. ulcerans* from the environment to humans ^3,15,17,21,22^. Our survey area was the Mornington Peninsula, where human Buruli ulcer cases have increased progressively over the past 20 years ^12^. Our key findings were that *M. ulcerans* was almost exclusively associated with one mosquito species in this region, *Ae. notoscriptus*, a mammalian host feeder, at a frequency of 0.87%. The estimated pathogen burden in each insect (Table S3) was consistent with the reported low infectious dose for *M. ulcerans* ^22,54^. Pathogen genomics provided support for a linked transmission chain between mosquitoes, native possums and humans in this region. The metabarcoding bloodmeal analysis of trapped mosquitoes revealed dual feeding by individual mosquitoes on both possums and humans, providing an example of a potential transmission pathway between infected possums and humans. Finally, spatial clustering analysis showed a striking overlap between clusters of possums shedding *M. ulcerans*, mosquitoes harbouring *M. ulcerans* and human Buruli ulcer cases, reinforcing the importance of possums and mosquitoes in the spread of *M. ulcerans* to humans.

Indications that invertebrates might be playing a role in the spread of Buruli ulcer first came from field surveys in West Africa in the late 1990s, where surveys in Benin identified water bugs in the genera *Naucoris* and *Diplonychus* as IS2404 PCR positive for *M. ulcerans* ^15,16,55^. Subsequent culture isolation of *M. ulcerans* from an aquatic insect in Benin (*Gerris* sp.) and potentially from *Naucoris* sp. in neighbouring Cote d’Ivoire further suggested a role for water bugs in disease transmission, at least in west Africa ^56,57^. A role for mosquitoes in the transmission of *M. ulcerans* anywhere was first revealed from entomological surveys in the mid-2000s on the Bellarine Peninsula and then more recently in Far North Queensland ^15,21,58^. Other observations that supported a role for mosquitoes in transmission included an analysis of the Buruli ulcer lesion location on the human body, which included more than 600 human Buruli ulcer cases, and showed a preponderance of lesions on ankles, elbows and backs of legs. These areas are frequently exposed and insects may preferentially bloodfeed there ^59^. Laboratory studies under one transmission scenario have also shown the competence of *Ae. notoscriptus* as mechanical vectors of *M. ulcerans* (Barnett criteria No. 4, Table 1) ^22^.

Of the ∼18,600 mosquitoes tested by IS2404 PCR from five different survey periods in the present study, *Ae. notoscriptus* was consistently positively associated with *M. ulcerans*. Where and how *M. ulcerans* is contaminating (or infecting) these mosquitoes remains to be determined. However, we made the somewhat unexpected observation that the most abundant mosquito species trapped on the Mornington Peninsula, *Cx. molestus,* which predominately feeds on avian species ^60^ was consistently IS2404 PCR negative. This observation might be explained by a difference in the ecology of these two mosquito species, such as how (or where) they are encountering *M. ulcerans.* The apparent *Ae. notoscriptus*/*M. ulcerans* tropism might also indicate a specific biological interaction between this insect and the mycobacterium, or alternatively, a *Cx. molestus* antagonism to mycobacterial carriage.

MLE is a parameter used in vector ecology to estimate the proportion of infected mosquitoes that are pathogen positive, based on a pooling-based screening assay, where the proportion is the parameter of a binomial distribution ^61^. The MLE of *Ae. notoscriptus* over all 5 years of our surveys was 5.88 *M. ulcerans* positive mosquitoes/1,000 tested (95% CI 4.37-7.76). The MLE for the only other positive mosquito species, *Ae. camptorhynchus* (consisting of two insects that tested positive for *M. ulcerans*), was 2.96/1,000 tested (95% CI 0.17-14.19). These estimates are consistent with those previously reported for *Ae. camptorhynchus* (10,558 mosquitoes tested, MLE 3.98 per 1,000) and *Ae. notoscriptus* (221 mosquitoes tested, MLE of 4.47 per 1,000) during the Bellarine Peninsula survey of that Buruli ulcer endemic area ^15^. Interestingly, these values are significantly higher than the MLE values from an *M. ulcerans* mosquito survey conducted in tropical far north Queensland, Australia which reported an MLE value of 0.13 /1,000 (95% CI 0.01–0.61) for different species of mosquito ^58^. To note though, there were few reported cases of Buruli ulcer in this area of Queensland during the field survey period, consistent with the suggestion that mosquito surveys for *M. ulcerans* will be useful for predicting Buruli ulcer risk in humans.

In tropical far north Queensland, Australia, host-seeking mosquitoes that harboured *M. ulcerans* collected through similar CO2 light traps to those used in this study were within the genera *Verallina*, *Coquillettidia* and *Mansonia* ^58^. These comparisons highlight that the mosquitoes encountering *M. ulcerans* appear to be region/environment specific, and these observations support the idea that mosquitoes are mechanical vectors of *M. ulcerans*. The major mosquito species associated with detection of *M. ulcerans* in Victoria, *Ae. notoscriptus, Ae. camptorhynchus*, *An. annulipes* and *Cq. linealis,* are all diverse opportunistic feeders.

To explore host sources for mosquitoes trapped in our study, we used metagenomic amplicon sequencing of engorged mosquitoes ^62–67^. Of the 90 blood-fed mosquitoes processed, 36 individual insects successfully had their host blood source identified. The 54 bloodmeals that were unidentified were probably due to degradation of the host DNA that occurs approximately 36 hours post-feeding ^68^. As the mosquitoes analysed in this study were collected from baited mosquito traps, the exact time post-feeding they were collected is unknown. Of the host blood sources identified, common ringtail possum was the most commonly identified bloodmeal, followed by birds and humans, with dual bloodmeals observed in 55% of mosquitoes (Figure 4, Figure S4). Most notably, three insects were determined to have bloodmeals from both common ringtail possums and humans. These insects consisted of two *Ae. notoscriptus* and one *Ae. rubrithorax*. Common ringtail possums are major wildlife reservoirs of *M. ulcerans* ^11^, and in this study we identified *M. ulcerans* in frequent association with *Ae. notoscriptus*. The identification of these dual bloodmeals provides evidence that *Ae. notoscriptus* mosquitoes are bloodfeeding between possums and humans within a relatively short timeframe. Dual bloodmeals may provide an opportunity for a mechanical transmission event of *M. ulcerans* from a possum with a Buruli ulcer teeming with *M. ulcerans,* from which a mosquito bloodfeeds and then moves to a nearby human. However, the observed positive association specifically between *Ae. notoscriptus* and *M. ulcerans* is not explained by our bloodmeal analysis, which showed several other mosquito species that were not *M. ulcerans* positive, such as *Cx. molestus*, *Ae. rubrithorax* and *Cq. Linealis,* also fed on possums. An alternative source of *M. ulcerans* acquisition for *Ae. notoscriptus* might be from possum excreta contaminating the *Ae. notoscriptus* breeding sites, as this species breeds in small artificial containers ^62^. Previous studies have shown that mosquito larvae can ingest *M. ulcerans* and test positive for *M. ulcerans* during the larval stage, but *M. ulcerans* is not detected in the pupa or adult, which is largely thought to be a result of the larval midgut being purged ^69^. However, if possum excreta, which can have a high concentration of *M. ulcerans* ^11^, falls into these artificial breeding sites, then this may provide a means to contaminate the adult *Ae. notoscriptus* during eclosion and emergence on top of the water’s surface.

In common with other arthropod-borne bacterial pathogens, *M. ulcerans* also has the genomic hallmarks of niche adaptation ^70,71^. For instance, the bacterial diseases vectored by blood-feeding arthropods, such as bubonic plague, spread to humans through the bite of fleas harbouring the bacterium *Yersinia pestis* ^72^, the tick-borne infections such as Lyme disease, Rocky Mountain Spotted Fever, ehrlichiosis (among others) caused by *Borrelia burgdorferi*, *Rickettsia rickettsii* and *Ehrlichia* sp., respectively ^73^, or tularemia, caused by infection with *Francisella tularensis* spread through the bite of infected ticks (with some subspecies also spread by mosquitoes) ^74–76^, are all caused by bacteria that have degenerating genomes. That is, like M. ulcerans, their genomes bear the distinctive hallmarks of evolutionary bottlenecks and niche-adaptation (plasmid acquisition, pseudogene accumulation, insertion sequence expansion), a genomic pattern thought indicative of a shared trajectory towards symbiosis with the arthropod host ^77^. In addition, the absence of recombination and highly clonal population structure of *M. ulcerans* is aligned with the population structure of bacteria in ‘closed symbiosis’ with their host animal ^78^.

Mosquitoes were not the only arthropods from which *M. ulcerans* were detected in the Mornington Peninsula. With over 20,500 arthropods collected using sticky traps (YST and SO), this study screened other insects that might broadly act as a mechanical vector, that is, an insect that is involved in the accidental transport of a pathogen ^79^ ^22^. *M. ulcerans* was successfully identified on two of ∼1,800 flies tested, identified as blowflies (*Calliphora hilli*). Previous studies have screened other species of flies, such as March flies (Tabanidae), with no positive detections ^58^. *C. hilli* is a native species occurring along the Australian east coast and in South Australia ^80^. This species is a carrion breeder occurring year-round throughout Victoria, with the female laying her eggs in decaying flesh where the larvae emerge ^80^. Possum carcases are readily infested by *C. hilli* ^81^, which may explain how this fly species is becoming contaminated with *M. ulcerans*, as possums are wildlife reservoirs of *M. ulcerans* ^10^. The likelihood of *C. hilli* carrying *M. ulcerans* to humans is relatively low due to this species rarely biting humans ^82^.

Interestingly, all species that have tested positive for *M. ulcerans* share quite different breeding environments and feeding preferences, although most feed on humans. *Ae. camptorhynchus* and *Verrallina sp.* are saltmarsh mosquitoes ^83,84^, while *Ae. notoscriptus* is a freshwater container breeder occurring in close association to humans ^85^. *Anopheles annulipes* and *Cx. australicus* are also freshwater breeding mosquitoes ^86^, although *Cx. australicus* typically feeds on avian species and is not considered to feed on humans ^87^. This diversity in mosquito species, habitats, flight range from breeding sites and host-preference feeding raises interesting questions about how these mosquitoes are coming into contact with *M. ulcerans* and how potential vector mosquito species may change based on the study area. Additionally, the ecology of the mosquito sub-species are generally poorly understood, including *Ae. Notoscriptus,* which occurs as a series of genetically differentiated clades within Australia ^39^.

In this study, we also deployed for the first time targeted enrichment genome capture using the Agilent RNA bait system to demonstrate beyond reasonable doubt the presence of *M. ulcerans* in association with mosquitoes ^52^. The IS2404, IS2606 and KR PCR assays have provided robust evidence for the presence of *M. ulcerans* in the environment, but detecting pathogen genomic DNA and using those sequences to forensically dissect transmission chains by matching pathogen genome SNP profiles is a powerful technological advance^52^. While the method lost sensitivity for IS2404 qPCR Ct values >32, we generated sufficient genome coverage from one IS2404 PCR positive mosquito and from two possum excreta specimens to show that the *M. ulcerans* genotypes of the captured genomes were identical to those associated with human Buruli ulcer cases in the study area, rather than Buruli ulcer cases linked to other regions (Figure 3). These findings provide strong support for mosquitoes and possums playing a role in the transmission of *M. ulcerans* in the Mornington Peninsula. These results also add to the growing body of evidence demonstrating the utility of pathogen genomics in specifically identifying the sources and transmission pathways of environmental pathogens and suggest that continued surveillance of *M. ulcerans* genotypes in mosquitoes and possums could be useful in guiding public health interventions that seek to control the spread of Buruli ulcer in the region. We are now exploring the use of genome sequence enrichment to dissect the genomic epidemiology from the perspective of environmental sources, in particular to track the spread of Buruli ulcer in Australian native possums across the region ^53^.

The SaTScan clustering analyses provide compelling evidence for the spatial association between the presence of *M. ulcerans* in possums, mosquitoes, and human Buruli ulcer cases in the study area (Figure 5). The triple-cluster overlap observed in our SaTScan analyses suggests that the spatial distribution of these three factors is likely to be closely linked and may be contributing to the persistence of Buruli ulcer in the Mornington Peninsula. Our finding of a low frequency of triple-cluster overlap among 10,000 randomisations, with only 8.9% of replicates showing this phenomenon, provides a level of confidence that our observations have not occurred randomly. This further supports the validity of our findings and reinforces the idea that the presence of *M. ulcerans* in possums and mosquitoes may play a crucial role in the transmission of *M. ulcerans* to humans in the study area. Our findings highlight the importance of ongoing surveillance of possum and mosquito populations, which may provide critical insights into the epidemiology of Buruli ulcer in the region and inform targeted public health interventions to control the disease.

Our research has some limitations. Although we found a spatial association between the presence of *M. ulcerans* in possums and mosquitoes, and human Buruli ulcer cases, we cannot absolutely conclude causality or directionality. In addition, the very specific set of circumstances that have led to the rise of Buruli ulcer in temperate southeastern Australia restricts the generalisability of our results. One must be cautious to draw parallels with African Buruli ulcer endemic countries for instance, where a highly susceptible mammalian reservoir equivalent to the Australian native possum is yet to be identified and evidence for mosquitoes as possible vectors is lacking ^88^. Further investigations are needed to better understand the underlying mechanisms that drive the association between the presence of *M. ulcerans* in these different species and the development of human Buruli ulcer cases.

Despite these caveats, our collective research over more than 15 years makes it very clear that mosquitoes are likely vectors and native possums are major wildlife reservoirs of *M. ulcerans* in southeastern Australia. Mosquito surveillance with *M. ulcerans* screening coupled with mosquito control and public health messaging to avoid mosquito bites are practical interventions that would be expected to reduce the incidence of human Buruli ulcer.

## Supporting information

Datafile S1

Datafile S2

Datafile S3

## Acknowledgments

We would like to acknowledge the residents of the Mornington Peninsula for assisting in collecting and sending sticky cards to our laboratory as part of our surveillance and to Mary Tachedjian, Michael Dunn, Simone Clayton, Julie Gaburro and Victoria Boyd for assistance during field work.

## Ethics

Ethical approval for the use in this study of de-identified human BU case location, aggregated at mesh block level, was obtained from the Victorian Government Department of Health Human Ethics Committee under HREC/54166/DHHS-2019-179235(v3), ‘Spatial risk map of Buruli ulcer infection in Victoria.’.

## Author Contributions

Conceptualisation PTM, JO, RF, SRC, PDRJ, JRW, AAH, KBG, TPS, SEL; Data curation PTM, KB, JO; Formal Analysis PTM, AHB, TPS; Funding acquisition SRC, PDRJ, KBG, JRW, AH, KG, TPS, SEL; Methodology PTM, AHB, KB, JCC, JRW, TPS, SEL; Writing – original draft PTM, AHB, SEL, TPS; Writing – review & editing all authors.

## Data availability statement

DNA sequences generated in this project are available under these GenBank accession numbers: PRJNA943595, OQ600123-4, OQ588831-67. Custom computer code: https://github.com/abuultjens/BU-3-way-SatScan; https://github.com/abuultjens/Mosquito_possum_human_genomic_analysis.

