## Supplementary data for "A transmission chain linking *Mycobacterium ulcerans* with *Aedes notoscriptus* mosquitoes, possums and human Buruli ulcer cases in southeastern Australia"

**List of supplementary material:**

**Tables**

Table S1: Accession details for DNA sequence reads generated and *M. ulcerans* genome sequence reads used in this study.
Table S2. Male mosquitoes collected and screened for *M. ulcerans* on the Mornington Peninsula.
Table S3. *M. ulcerans* qPCR testing results for positive IS2404, IS2606 and KR in *Ae. notoscriptus, Ae. camptorhynchus* and *C. hilli*.
Table S4. Breakdown of the number of individual or pools of female *Ae. notoscriptus* screened each year.

**Figures**

Figure S1. Comparison of the size of the qPCR amplicon run on a 1% agarose gel between a real *M. ulcerans* positive control and the synthetic positive control which was used for the screening assays
Figure S2. IS2404 qPCR standard curve
Maximum likelihood phylogenetic tree based on an 874 bp region of the COI gene
Figure S3. SNP imputation validation.
Figure S4: Tanglegrams analysis
Figure S5. Maximum likelihood phylogenetic tree based on an 871 bp region of the COI gene.
Figure S6. Mosquito bloodmeal analysis all detected bloodmeal sources.

**Datasets**

Datafile S1: Oligonucleotide sequences of RNA baits used for *M. ulcerans* genome-enrichment sequencing.
Datafile S2: Mosquito trapping details
Datafile S3: *M. ulcerans* SNP alleles used for phylogenomic inference

**Table S1.** *Mycobacterium ulcerans* genome sequence reads used in this study

| Isolate ID | Alternate ID | Genbank accession | Year | Location | Region | Host species | Sequencing chemistry | Sequencing platform | Country | Reference |
|---|---|---|---|---|---|---|---|---|---|---|
| DMG2212098-DMG2301551 | 5675 | PRJNA943595 | 2019 | Victoria, Australia | South Mornington Peninsula | *Aedes notoscriptus* | NextSeq 2x150bp | Illumina MiSeq and NextSeq 2000 | Australia | This study |
| DMG2212099 | 4601-02 | PRJNA943595 | 2019 | Victoria, Australia | South Mornington Peninsula | *Aedes notoscriptus* | MiSeq 2x150bp | Illumina MiSeq | Australia | This study |
| DMG2212100 | 4997 | PRJNA943595 | 2019 | Victoria, Australia | South Mornington Peninsula | *Aedes notoscriptus* | MiSeq 2x150bp | Illumina MiSeq | Australia | This study |
| DMG2304587 | 226 | PRJNA943595 | 2021 | Victoria, Australia | South Mornington Peninsula | *Aedes notoscriptus* | NextSeq 2x150bp | NextSeq 2000 | Australia | This study |
| DMG2304588 | 819 | PRJNA943595 | 2021 | Victoria, Australia | South Mornington Peninsula | *Aedes notoscriptus* | NextSeq 2x150bp | NextSeq 2000 | Australia | This study |
| DMG2300866 | RES21-00999 | PRJNA943595 | 2022 | Victoria, Australia | South Mornington Peninsula | Common ringtail possum feces | NextSeq 2x150bp | NextSeq 2000 | Australia | This study |
| DMG2300867 | RES21-00921 | PRJNA943595 | 2022 | Victoria, Australia | South Mornington Peninsula | Common ringtail possum feces | NextSeq 2x150bp | NextSeq 2000 | Australia | This study |
| DMG1701366 | | SRR6346235 | 2015 | Victoria, Australia | South Mornington Peninsula | Human | NextSeq 2x150bp | NextSeq 500 | Australia | (1) |
| DMG1701349 | | SRR6346254 | 2015 | Victoria, Australia | South Mornington Peninsula | Human | NextSeq 2x150bp | NextSeq 500 | Australia | (1) |
| DMG1701351 | | SRR6346282 | 2015 | Victoria, Australia | North Mornington Peninsula | Human | NextSeq 2x150bp | NextSeq 500 | Australia | (1) |
| DMG1701353 | | SRR6346280 | 2015 | Victoria, Australia | South Mornington Peninsula | Human | NextSeq 2x150bp | NextSeq 500 | Australia | (1) |
| DMG1701357 | | SRR6346246 | 2015 | Victoria, Australia | South Mornington Peninsula | Human | NextSeq 2x150bp | NextSeq 500 | Australia | (1) |
| DMG1701358 | | SRR6346247 | 2015 | Victoria, Australia | South Mornington Peninsula | Human | NextSeq 2x150bp | NextSeq 500 | Australia | (1) |
| DMG1701359 | | SRR6346240 | 2015 | Victoria, Australia | Bellarine Peninsula | Human | NextSeq 2x150bp | NextSeq 500 | Australia | (1) |
| DMG1701360 | | SRR6346241 | 2015 | Victoria, Australia | South Mornington Peninsula | Human | NextSeq 2x150bp | NextSeq 500 | Australia | (1) |
| DMG1701361 | | SRR6346238 | 2015 | Victoria, Australia | South Mornington Peninsula | Human | NextSeq 2x150bp | NextSeq 500 | Australia | (1) |
| DMG1701362 | | SRR6346239 | 2015 | Victoria, Australia | South Mornington Peninsula | Human | NextSeq 2x150bp | NextSeq 500 | Australia | (1) |
| DMG1701365 | | SRR6346234 | 2015 | Victoria, Australia | Bellarine Peninsula | Human | NextSeq 2x150bp | NextSeq 500 | Australia | (1) |
| DMG1701368 | | SRR6346243 | 2015 | Victoria, Australia | Bellarine Peninsula | Human | NextSeq 2x150bp | NextSeq 500 | Australia | (1) |
| DMG1701371 | | SRR6346364 | 2016 | Victoria, Australia | Bellarine Peninsula | Human | NextSeq 2x150bp | NextSeq 500 | Australia | (1) |
| DMG1701373 | | SRR6346362 | 2016 | Victoria, Australia | North Mornington Peninsula | Human | NextSeq 2x150bp | NextSeq 500 | Australia | (1) |

| | | | | | | | | | | |
|---|---|---|---|---|---|---|---|---|---|---|
| DMG1701376 | | SRR6346359 | 2016 | Victoria, Australia | Bellarine Peninsula | Human | NextSeq 2x150bp | NextSeq 500 | Australia | (1) |
| DMG1701377 | | SRR6346358 | 2016 | Victoria, Australia | South Mornington Peninsula | Human | NextSeq 2x150bp | NextSeq 500 | Australia | (1) |
| DMG1701384 | | SRR6346374 | 2016 | Victoria, Australia | Bellarine Peninsula | Human | NextSeq 2x150bp | NextSeq 500 | Australia | (1) |
| DMG1701389 | | SRR6346299 | 2016 | Victoria, Australia | North Mornington Peninsula | Human | NextSeq 2x150bp | NextSeq 500 | Australia | (1) |
| DMG1701393 | | SRR6346303 | 2016 | Victoria, Australia | North Mornington Peninsula | Human | NextSeq 2x150bp | NextSeq 500 | Australia | (1) |
| DMG1701396 | | SRR6346304 | 2005 | Victoria, Australia | North Mornington Peninsula | Human | MiSeq 2x300bp | Illumina MiSeq | Australia | (1) |
| DMG1701397 | | SRR6346297 | 2005 | Victoria, Australia | Gippsland | Human | MiSeq 2x300bp | Illumina MiSeq | Australia | (1) |
| DMG1701398 | | SRR6346296 | 2006 | Victoria, Australia | Bellarine Peninsula | Human | MiSeq 2x150bp | NextSeq 500 | Australia | (1) |
| DMG1701400 | | SRR6346331 | 2015 | Victoria, Australia | North Mornington Peninsula | Human | NextSeq 2x150bp | NextSeq 500 | Australia | (1) |
| DMG1701403 | | SRR6346330 | 2007 | Victoria, Australia | North Mornington Peninsula | Human | MiSeq 2x150bp | NextSeq 500 | Australia | (1) |
| DMG1701404 | | SRR6346335 | 2008 | Victoria, Australia | North Mornington Peninsula | Human | MiSeq 2x150bp | NextSeq 500 | Australia | (1) |
| DMG1701405 | | SRR6346336 | 2008 | Victoria, Australia | Bellarine Peninsula | Human | MiSeq 2x150bp | NextSeq 500 | Australia | (1) |
| DMG1701406 | | SRR6346333 | 2008 | Victoria, Australia | Bellarine Peninsula | Dog | MiSeq 2x150bp | NextSeq 500 | Australia | (1) |
| DMG1701409 | | SRR6346328 | 2008 | Victoria, Australia | Bellarine Peninsula | Human | MiSeq 2x150bp | NextSeq 500 | Australia | (1) |
| DMG1701420 | | SRR6346276 | 2009 | Victoria, Australia | Gippsland | Koala | MiSeq 2x150bp | NextSeq 500 | Australia | (1) |
| DMG1701423 | | SRR6346279 | 2009 | Victoria, Australia | North Mornington Peninsula | Human | MiSeq 2x150bp | NextSeq 500 | Australia | (1) |
| DMG1701428 | | SRR6346292 | 2002 | Victoria, Australia | Gippsland | Human | MiSeq 2x300bp | Illumina MiSeq | Australia | (1) |
| DMG1701435 | | SRR6346206 | 2002 | Victoria, Australia | Gippsland | Long Footed Potoroo | MiSeq 2x300bp | Illumina MiSeq | Australia | (1) |
| DMG1701442 | | SRR6346230 | 2002 | Victoria, Australia | Gippsland | Human | MiSeq 2x300bp | Illumina MiSeq | Australia | (1) |
| DMG1701477 | | SRR6346351 | 2003 | Victoria, Australia | Gippsland | Human | MiSeq 2x300bp | Illumina MiSeq | Australia | (1) |
| DMG1701493 | | SRR6346340 | 2015 | Victoria, Australia | Gippsland | Human | NextSeq 2x150bp | NextSeq 500 | Australia | (1) |
| DMG1701495 | | SRR6346339 | 2005 | Victoria, Australia | Gippsland | Human | MiSeq 2x300bp | Illumina MiSeq | Australia | (1) |

**Table S2.** Male mosquitoes collected and screened for *M. ulcerans* on the Mornington Peninsula.

| Species | No. collected/No. screened (No. positive for *M. ulcerans*) | | | | | Total |
|---|---|---|---|---|---|---|
| | Dec 16-Apr 17 | Nov 17-May 18 | Dec 18-May 19 | Nov 19-Mar 20 | Feb 21-Mar 21 | |
| *Ae. notoscriptus* | | 2/2 | 7/7 | 44/24 | 5/5 | 58/38 |
| *Ae. rubrithorax* | | | 1/1 | | | 1/1 |
| *Cq. linealis* | 9/9 | 24/24 | | 67/17 | | 100/50 |
| *Cx. australicus* | | 11/11 | 2/2 | 303/16 | | 316/29 |
| *Cx. globocoxitus* | | | | 619/175 | | 619/175 |
| *Cx. molestus* | 1/1 | | 7/7 | 69/14 | 2/2 | 79/24 |
| *Cx. quinquefasciatus* | | 32/32 | 1/1 | 108/76 | | 141/109 |
| *Tp. tasmaniensis* | | | | 3/3 | | 3/3 |
| **Total** | **10/10** | **69/69** | **18/18** | **1,213/325** | **7/7** | 1,317/429 |

**Table S3.** *Mycobacterium ulcerans* qPCR testing results for positive IS2404, IS2606 and KR in *Ae. notoscriptus*, *Ae. camptorhynchus* and *C. hilli*.

| Species (season collected) | Location | qPCR assay | | | *M. ulcerans* genome equivalents/mosquito[1] |
|---|---|---|---|---|---|
| | | IS2404 | IS2606 | KR | |
| *Ae. notoscriptus* (16/17) | Sorrento | 33.64 | 35.18 | 34.94 | 401 |
| *Ae. notoscriptus* (16/17) | Tootgarook | 36.41 | - | - | 76 |
| *Ae. notoscriptus* (17/18) | Capel Sound | 38.21 | - | - | 26 |
| *Ae. notoscriptus* (17/18) | Capel Sound | 36.34 | NT[2] | - | 79 |
| *Ae. notoscriptus* (17/18) | Capel Sound | 38.27 | - | - | 25 |
| *Ae. notoscriptus* (17/18) | Capel Sound | 36.89 | - | - | 57 |
| *Ae. notoscriptus* (17/18) | St Andrews Beach | 35.02 | - | - | 175 |
| *Ae. notoscriptus* (17/18) | Tootgarook | 32.62 | NT | 32.18 | 740 |
| *Ae. notoscriptus* (17/18) | Tootgarook | 38.22 | NT | - | 25 |
| *Ae. notoscriptus* (17/18) | Tootgarook | 33.40 | - | 34.76 | 463 |
| *Ae. notoscriptus* (18/19) | Rye | 36.79 | NT | 32.31 | 60 |
| *Ae. notoscriptus* (18/19) | Rye | 36.67 | NT | 35.91 | 65 |
| *Ae. notoscriptus* (18/19) | Rye | 36.06 | NT | - | 93 |
| *Ae. notoscriptus* (18/19) | Rye | 35.56 | 39.371 | 34.95 | 126 |
| *Ae. notoscriptus* (18/19) | Rye | 36.34 | NT | 35.61 | 79 |
| *Ae. notoscriptus* (18/19) | Rye | 36.07 | 38.01 | 35.41 | 93 |
| *Ae. notoscriptus* (18/19) | Rye | 33.70 | 32.49 | 31.62 | 386 |
| *Ae. notoscriptus* (18/19) | Rye | 29.74 | 38.36 | 33.36 | 4183 |
| *Ae. notoscriptus* (18/19) | Rye | 31.51 | 38.90 | 33.67 | 1443 |
| *Ae. notoscriptus* (18/19) | Rye | 36.89 | NT | - | 57 |
| *Ae. notoscriptus* (18/19) | Rye | 33.32 | 39.00 | 35.92 | 486 |
| *Ae. notoscriptus* (18/19) | Rye | 37.25 | - | 35.80 | 46 |
| *Ae. notoscriptus* (18/19) | Rye | 35.11 | - | 34.19 | 165 |
| *Ae. notoscriptus* (18/19) | Rye | 36.44 | 40.07 | 35.02 | 74 |
| *Ae. notoscriptus* (18/19) | Rye | 35.27 | 45.00 | 35.49 | 150 |
| *Ae. notoscriptus* (19/20) | Blairgowrie | 35.60 | 37.90 | 33.02 | 123 |
| *Ae. notoscriptus* (19/20) | Blairgowrie | 38.56 | 39.89 | - | 21 |
| *Ae. notoscriptus* (19/20) | Blairgowrie | 37.45 | 38.58 | 35.23 | 40 |
| *Ae. notoscriptus* (19/20) | Rye | 38.11 | 39.96 | 38.86 | 27 |
| *Ae. notoscriptus* (19/20) | Rye | 35.14 | 39.55 | - | 163 |
| *Ae. notoscriptus* (19/20) | Rye | 39.15 | - | - | 15 |
| *Ae. notoscriptus* (19/20) | Rye | 33.02 | 35.18 | 32.44 | 582 |
| *Ae. notoscriptus* (19/20) | Rye | 36.76 | 37.03 | 35.15 | 61 |
| *Ae. notoscriptus* (19/20) | Rye | 37.96 | 38.67 | 33.48 | 30 |
| *Ae. notoscriptus* (19/20) | Rye | 36.09 | 34.62 | 32.86 | 92 |
| *Ae. notoscriptus* (19/20) | Rye | 37.86 | 39.25 | 39.26 | 32 |
| *Ae. notoscriptus* (19/20) | Rye | 36.51 | 36.79 | 34.07 | 71 |
| *Ae. notoscriptus* (19/20) | Sorrento | 37.73 | 37.08 | 34.86 | 34 |
| *Ae. notoscriptus* (19/20) | Sorrento | 38.51 | 38.80 | 36.34 | 21 |
| *Ae. notoscriptus* (20/21) | Blairgowrie | 33.47 | 34.52 | 29.36 | 489 |
| *Ae. notoscriptus* (20/21) | Blairgowrie | 34.15 | 35.05 | 30.10 | 295 |
| *Ae. notoscriptus* (20/21) | Blairgowrie | 38.64 | 40.21 | 35.24 | 20 |
| *Ae. notoscriptus* (20/21) | Blairgowrie | 31.20 | 33.26 | 28.52 | 1738 |
| *Ae. notoscriptus* (20/21) | Blairgowrie | 39.65 | - | - | 11 |
| *Ae. notoscriptus* (20/21) | Rye | 37.50 | - | NT | 39 |
| *Ae. notoscriptus* (20/21) | Rye | 37.48 | - | - | 40 |
| *Ae. camptorhynchus* (19/20) | Sorrento | 41.21 | 43.10 | 35.25 | |
| *C. hilli* (18/19) | Rye | 35.9215 | 34.69 | - | |
| *C. hilli* (18/19) | Rye | 37.541 | - | 38.26 | |

[1] Note that insect DNA was eluted in 100uL and 2uL was used for qPCR; [2]NT = Not tested

**Table S4.** Breakdown of the number of individual or pools of female *Ae. notoscriptus* that were screened each year.

| Year | Screening size | No. individuals or pools | Number positive |
|---|---|---|---|
| 2016-17 | Individuals screened | 4 | 1 |
| | Pools <15 (total no *Ae. notoscriptus* screened in pools 170) | 23 | 1 |
| 2017-18 | Individuals screened | 367 | 8 |
| | Pools <15 | 0 | 0 |
| 2018-19 | Individuals screened | 112 | 3 |
| | Pools <15 (total no *Ae. notoscriptus* screened in pools 1667) | 341 | 12 |
| 2019-20 | Individuals screened | 448 | 3 |
| | Pools <15 (total no *Ae. notoscriptus* screened in pools 3882) | 383 | 11 |
| 2021 | Individuals screened | 1247 | 7 |
| | Pools <15 | 0 | 0 |

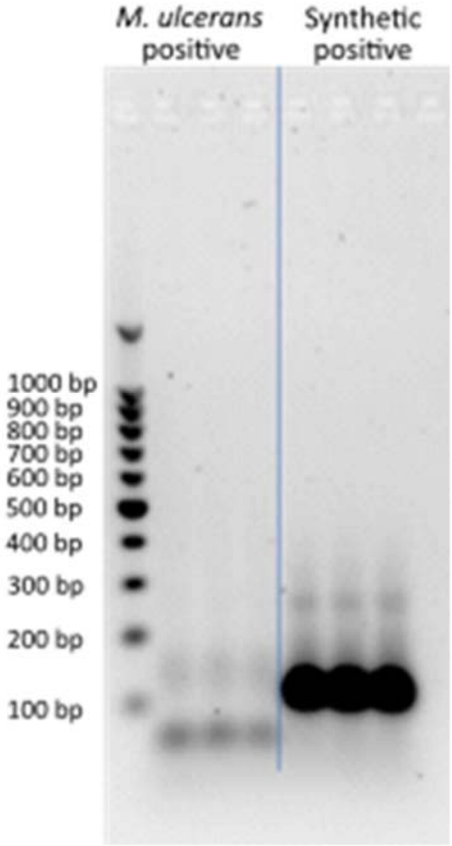

***Figure S1.*** *Comparison of the size of the qPCR amplicon run on a 1% agarose gel between a real M. ulcerans positive control and the synthetic positive control which was used for the screening assays.*

A

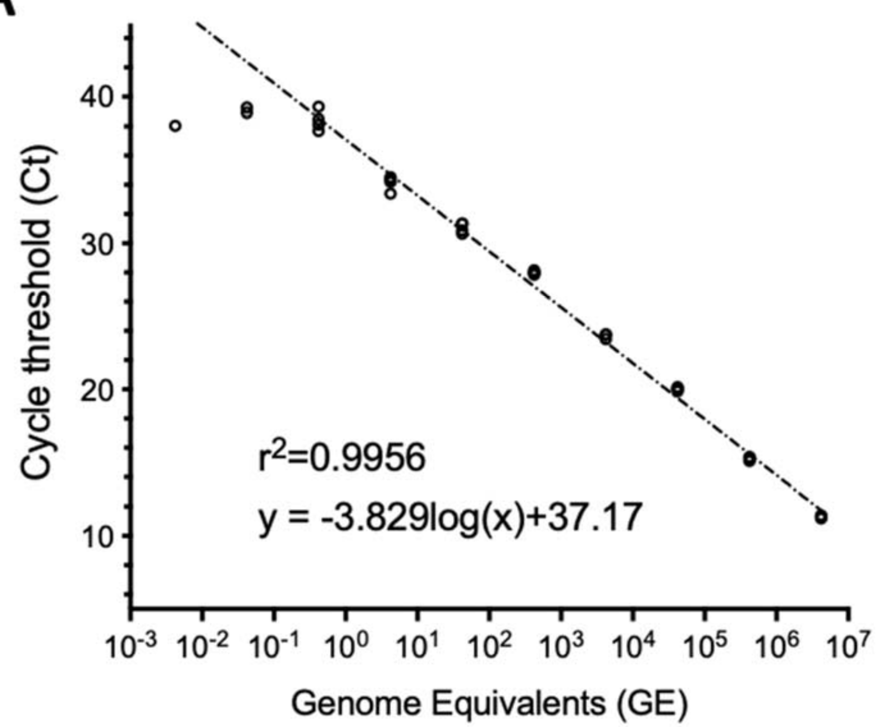

B

| gDNA conc. (ng/uL) | Genome Equivalents | Rep 1 | Rep 2 | Rep 3 | Rep 4 |
|---|---|---|---|---|---|
| **12.1** | 4200000 | 11.34 | 11.2 | 11.4 | 11.27 |
| **1.2** | 420000 | 15.09 | 15.17 | 15.35 | 15.35 |
| **0.12** | 42000 | 19.87 | 20.02 | 20.05 | 20.15 |
| **0.012** | 4200 | 23.75 | 23.74 | 23.44 | - |
| **0.0012** | 420 | 27.95 | 28.02 | 27.84 | 28.16 |
| **0.00012** | 42 | 30.89 | 31.34 | 31.35 | 30.66 |
| **0.000012** | 4.2 | 34.38 | 33.4 | 34.53 | 34.21 |
| **0.0000012** | 0.42 | 39.32 | 37.68 | 38.47 | 38.12 |

***Figure S2:*** *IS2404 qPCR standard curve. (A) Plot of 10-fold dilution series of M. ulcerans genomic DNA tested by IS2404 Taqman real-time PCR (each dilution tested in quadruplicate). The x-axis is $log_{10}$ scale. (B) Cycle-threshold (Ct) values for each genomic DNA dilution used to prepare the plot in (A).*

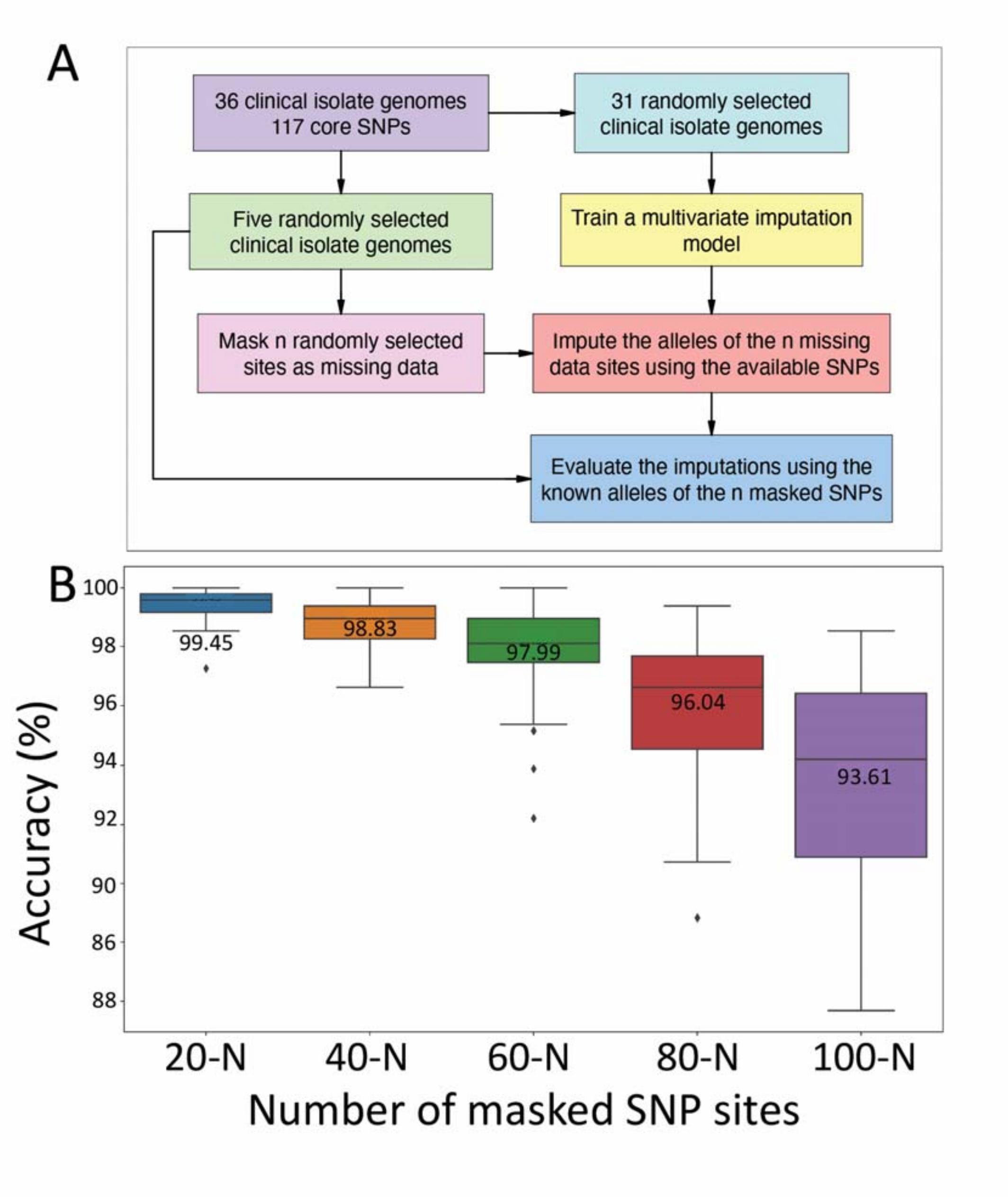

***Figure S3:*** *Validation of allele imputation approach to assess robustness to level of missing data and genome selection. (A) Flow diagram of imputation method. (B) Boxplots showing the distribution of imputation accuracy across 100 replicates for varying numbers of M. ulcerans chromosome SNP sites masked as missing data for five randomly selected M. ulcerans clinical isolate genomes. The line inside each box represents the median accuracy and the annotated numbers above each box indicate the mean accuracy in percentage. The whiskers extend to 1.5x the interquartile range.*

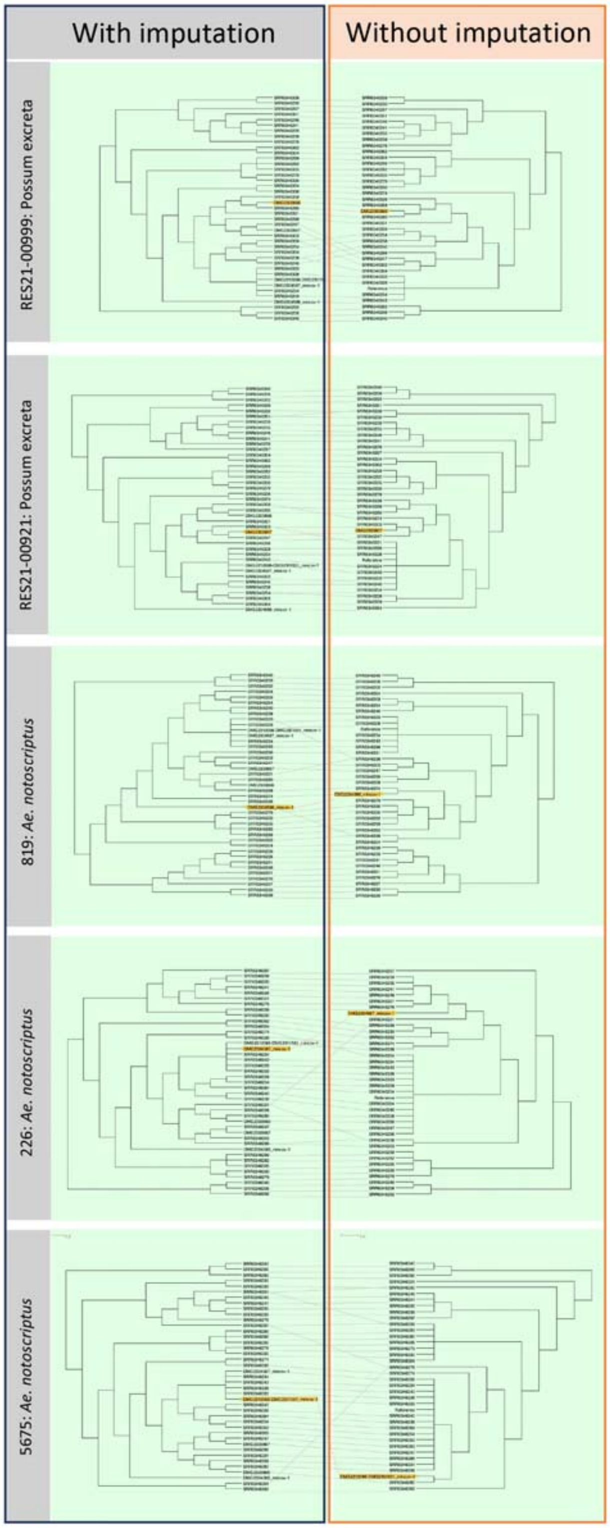

***Figure S4:*** *Tanglegrams analysis showing impact of imputation on phylogenetic inference. Depicted are tanglegrams (Dendroscope) showing phylogenies inferred* *<u>with</u>* *imputed SNP alignments containing each of the five, individual sequence enrichment and 36 clinical isolate genome sequences compared to phylogenies of the individual sequence enrichment genomes and*

*36 clinical isolates without imputation. Yellow highlighted tips show the sequence enrichment genomes. Branch lengths transformed and presented as cladograms.*

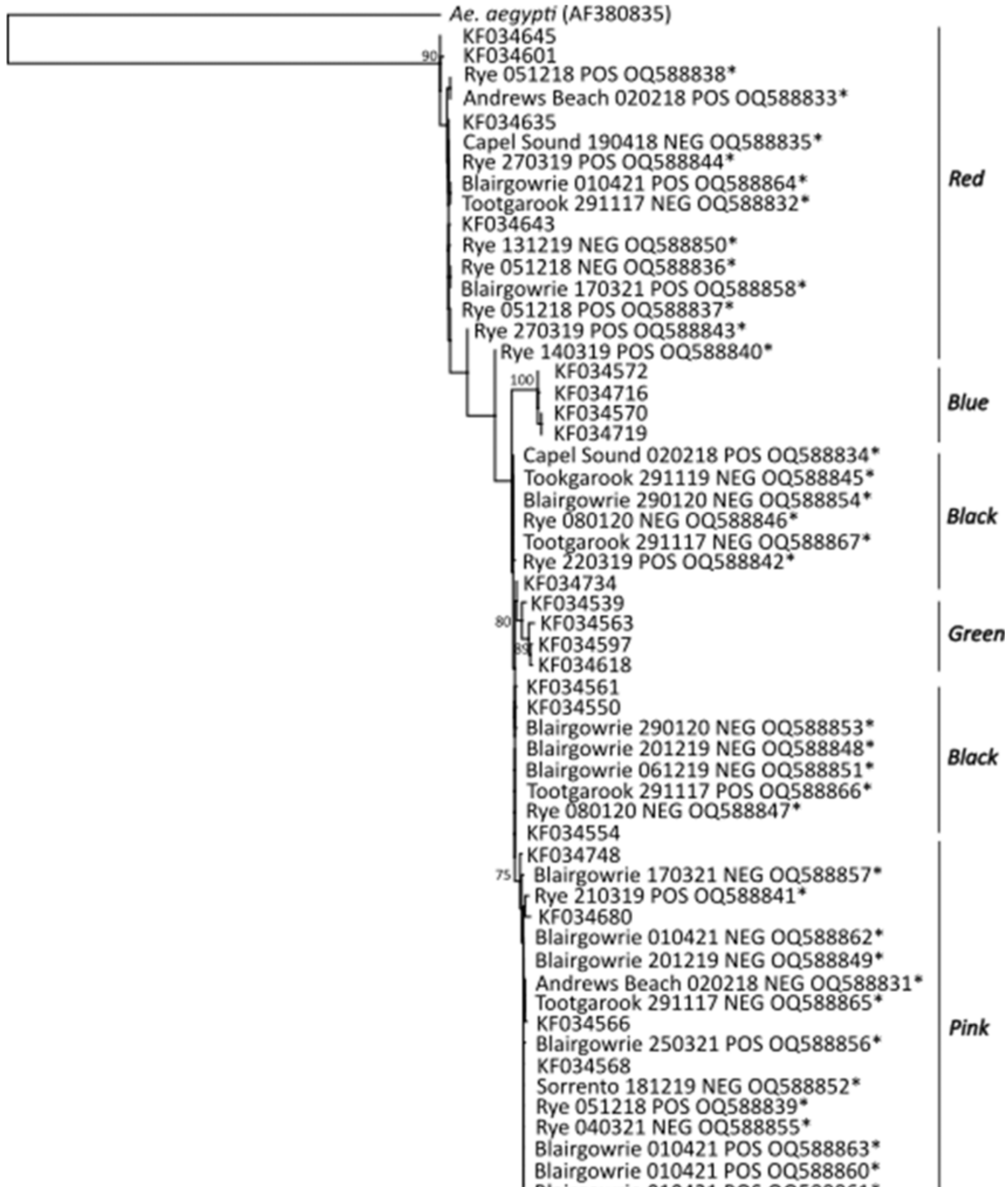

***Figure S5. Maximum likelihood phylogenetic tree based on an 871 bp region of the COI gene****.* *Labelling indicates capture location, collection date, M. ulcerans infection status and accession number. Clades are labelled based on Endersby et al. 2013, with four well-supported clades (blue, red, green and pink) and a fifth poorly supported clade (black). Insects from the blue clade had previously been collected in the subtropics of Queensland (QLD), green in the tropics of QLD, pink in temperate southern Australia, red from a broad distribution along the east coast and black from New Zealand and south-eastern Australia. The Hasegawa-Kishino-Yano (HKY) substitution model was used with 1,000 bootstrap replicates. Bootstrap proportions (BSP ≥ 70%) are indicated alongside the nodes. The number of nucleotide substitutions per site is represented by the scale bar. Aedes aegypti was used as an outgroup. * sequences generated in this study.*
